# Genome and transcriptome architecture of allopolyploid okra *(Abelmoschus esculentus)*

**DOI:** 10.1101/2021.11.18.469076

**Authors:** Ronald Nieuwenhuis, Thamara Hesselink, Hetty C. van den Broeck, Jan Cordewener, Elio Schijlen, Linda Bakker, Sara Diaz Trivino, Darush Struss, Simon-Jan de Hoop, Hans de Jong, Sander A. Peters

## Abstract

We present the first annotated genome assembly of the allopolyploid okra (*Abelmoschus esculentus*). Analysis of telomeric repeats and gene rich regions suggested we obtained whole chromosome and chromosomal arm scaffolds. Besides long distal blocks we also detected short interstitial TTTAGGG telomeric repeats, possibly representing hallmarks of chromosomal speciation upon polyploidization of okra. Ribosomal RNA genes are organized in 5S clusters separated from the 18S-5.8S-28S units, clearly indicating an S-type rRNA gene arrangement. The assembly is consistent with cytogenetic and cytometry observations, identifying 65 chromosomes and 1.45Gb of expected genome size in a haploid sibling. Approximately 57% of the genome consists of repetitive sequence. BUSCO scores and A50 plot statistics indicated a nearly complete genome. Kmer distribution analysis suggests that approximately 75% has a diploid nature, and at least 15% of the genome is heterozygous. We did not observe aberrant meiotic configurations, suggesting there is no recombination among the sub-genomes. BUSCO configurations as well as k-mer clustering analysis pointed to the presence of at least 2 sub-genomes. These observations are indicative for an allopolyploid nature of the okra genome. Structural annotation, using gene models derived from mapped IsoSeq transcriptome data, generated over 130,000 putative genes. Mapped transcriptome data from public okra accessions of Asian origin confirmed the predicted genes, showing limited genetic diversity of 1SNP/2.1kb. The discovered genes appeared to be located predominantly at the distal ends of scaffolds, gradually decreasing in abundance toward more centrally positioned scaffold domains. In contrast, LTR retrotransposons were more abundant in centrally located scaffold domains, while less frequently represented in the distal ends. This gene and LTR-retrotransposon distribution is consistent with the observed heterochromatin organization of pericentromeric heterochromatin and distal euchromatin. The derived amino acid queries of putative genes were subsequently used for phenol biosynthesis pathway annotation in okra. Comparison against manually curated reference KEGG pathways from related *Malvaceae* species revealed the genetic basis for putative enzyme coding genes that likely enable metabolic reactions involved in the biosynthesis of dietary and therapeutic compounds in okra.

## Introduction

The well-known okra (*Abelmoschus esculentus*) vegetable belongs to the family *Malvaceae*, comprising more than 244 genera and over 4,200 species. The *Malvaceae* are divided into 9 subfamilies of which okra belongs to the subfamily *Malvoideae. Abelmoschus* is closely related to *Hibiscus* species like *Hibiscus rosa-chinenesis* or Chinese rose and *Hibiscus cannabinus* or Kenaf, which is exemplified by the beautiful characteristic Hibiscus-like flowers that both genera display. Based on genetic differences, *Abelmoschus* has now been placed in a separate genus from *Hibiscus* though. Okra is flowering continuously and is self-compatible, however cross-pollination up to 20% has been reported. Its characteristic hermaphroditic flowers usually have white or yellow perianths, consisting of five petals and five sepals, whereas calyx, corolla and stamens are fused at the base. Other well-known species in the *Malvaceae* are cocoa (*Theobroma cacao*), cotton (*Gossypium hirsutum*), and *Tilia* species like lime tree, the mangrove *Heritiera* species and durian (*Durio zibethinus*). The genus *Abelmoschus* contains 11 species, four subspecies and five varieties (Li *et al*., 2020) of which most members have economic value. Okra or ‘lady’s finger’ is a low-calorie vegetable, mainly cultivated for its fruits that are harvested while still unripe, containing a large variety of nutrients and elements essential for daily human consumption, such as vitamins, flavonoids, minerals, and other health components such as folate and fibers (Muimba-Kankolonga, 2018; Wu *et al*., 2020). For example, total polyphenol extracts from okra fruits, containing flavonoids such as myricitin and quercitin, have been demonstrated for their antidiabetic activity in obese rats suffering from type 2 diabetes mellitus (Peter *et al*., 2021). These health compounds and additional nutritional qualities make okra an appreciated vegetable in many parts of the tropics and subtropics of Asia, Africa and America, gaining rapidly in popularity. Global production has increased yearly since 1994, reaching 10M tonnes in 2019 and covering some 2.5M ha (http://faostat.fao.org), with Asia having the largest production share of almost 70%, of which India alone is currently annually producing more than 4M tonnes. However, its production is challenged by a range of pathogens and insect pests, such as powdery mildew and blackmold (*Cerospora abelmoschii*), bacterial blight disease (*Xanthomonas campestris* p.v. *malvacearum*), mycoplasmas, nematodes, worms and insects such as whitefly (*Bemisia tabaci*), thrips (*Thrips palmi*), cotton leafhopper (*Amarasca biguttula*) and aphids (*Aphis gossypii*). Besides feeding damage, these vectors can transmit viruses such as Yellow Vein Mosaic Virus (YVMV), a geminivirus, causing crop losses of up to 80-90% without pest control (Benchasri, 2012; Muimba-Kankolonga, 2018; Dhankhar and Koundinya, 2020; Lata *et al*., 2021). Typical symptoms of YVMV infected okra plants are a stunted growth, with vein and veinlets turning yellow in colour, producing seed pods that are small, distorted, and chlorotic. Crop loss may be reduced to 20-30%, by controlling insect pests with rather harmful and toxic chemicals and insecticides (Ali *et al*., 2005), causing considerable collateral damage to the ecosystem. Moreover, increased insect tolerance to pesticides has led to over-use and mis-use of chemicals, leaving unhealthy high levels of pesticide residues (Benchasri, 2012). Although there are some YVMV tolerant okra genotypes, such as Nun1144 and Nun1145 (Venkataravanappa *et al*., 2013), the genetic basis for this tolerance has not been identified. Besides a need for disease resistance, other breeding challenges and demands include maximizing production, unravelling the genetic basis for abiotic stress tolerance, and the need to develop double haploid lines enabling the study of recessive gene traits (Dhankhar and Koundinya, 2020).

To meet current demands and challenges, accelerated breeding is urgently needed. Presently, several methods of breeding for improvement in okra are being used, such as pure line selection, pedigree breeding, as well as mutation and heterosis breeding (Dhankhar and Koundinya, 2020). These methods are very time consuming though, and often involve laborious analyses over multiple generations. Despite wide genetic variation available among wild relatives of okra, significant crop improvement by introgression breeding, has not been achieved due to hybridization barriers. Advanced breeding is further hampered due to the lack of sufficient molecular markers, linkage maps and reference genome, and this in turn has impeded genome and transcriptome studies. Molecular studies have further been complicated due to the presence of large amounts of mucilaginous and polyphenolic compounds in different tissues, interfering with the preparation of genetic materials (Takakura and Nishio, 2012; Lata *et al*., 2021). Furthermore, correct *de novo* assembly is presumed to be complex because of the expected large genome and transcriptome size and the highly polyploid nature of the genome. Salameh (2014) reported flow-cytometric estimates of nuclear DNA size estimations with 2C values ranging from 3.98 to 17.67 pg, equaling to genome sizes between 3.8 to 17.3 Gbp. In addition, chromosome counts demonstrated a huge variation, ranging from 2n=62 to 2n=144, with 2n=130 as the most frequently observed chromosome number (Benchasri, 2012; Merita *et al*., 2012). These findings have led to further assumptions on the geographical origin of cultivated *A. esculentus*, speculating that a 2n=58 specie *A. tuberculatus* native from Northern India and a 2n=72 specie *A. ficulneus* from East Africa might have hybridized followed by a chromosome doubling, giving rise to an alloploid *Abelmoschus* hybrid with 2n=130 (Joshi and Hardas, 1956; Siemonsma, 1982; Benchasri, 2012; Merita *et al*., 2012). However, genomic, genetic and cytological information is scanty, limiting the possibilities to further understand the hereditary constituent of the crop. In this study we benefitted from naturally occurring okra haploids, circumventing heterozygosity in the reconstruction of composite genome sequences, supporting faithful genome reconstruction (Langley *et al*., 2011). Here we present a detailed insight into the complex genome and transcriptome architecture of an okra cultivar and its haploid descendent, using cytogenetic characterization of its mitotic cell complements and meiosis, and advanced sequencing and assembly technologies of the haploid genome, providing basic scientific knowledge for further evolutionary studies and representing a necessary resource for future molecular based okra breeding. Furthermore, we provide a structural and functional genome annotation that is of paramount importance to understand plant metabolism (Weissenborn *et al*., 2017) and the genetic basis for the enzyme coding genes that enable metabolic reactions involved in the biosynthesis of dietary and therapeutic compounds in okra.

## Results and discussion

### Cytogenetic characterization of the okra crop

As okra is known to contain large numbers of chromosomes that differ between cultivars, we first established chromosome counts and morphology in the cultivar used in this study. Actively growing root tips were fixed and prepared for cell spread preparations following a standard pectolytic enzyme digestion and airdrying protocol, and DAPI fluorescence microscopy (Kantama *et al*., 2017). In this diploid red petiole phenotype plants, we counted 130 chromosomes in late prophase and metaphase cell complements (Figures 1a). Chromosomes measured 1-2 µm, often show telomere to telomere interconnections (Figure 1b) and were clearly monocentric (Figure 1a, arrows). In addition, a few chromosomes displayed a less condensed distal region at one of the chromosome arms and satellites (Figure 1c,d), which we interpreted as the nucleolar organiser region (NOR) of the satellite chromosome. Interphase nuclei showed well differentiated heterochromatic domains or chromocenters, most of them with more than 130 spots, although a small number of nuclei decondensed most of its heterochromatin, leaving only a striking pattern of about 10 chromocenters (Figure 1e). We next applied flow cytometry on DAPI stained nuclei, using young leaf material from five normally growing red petiole phenotype plantlets. Since we expected a considerable DNA content for okra nuclei, we decided to use a reference sample from Agave (*Agave americana*), which has a known DNA content of 15.90 picogram. Surprisingly, in comparison to the Agave reference flow cytometric profile, the 2C DNA amount for the normal okra plant was estimated at 2.99 pg ±0.01 (Table S1). This amount is equivalent to a genome size of approximately 2.92 Mbp. In contrast to 130 chromosomes for the diploid okra, we observed a chromosome number of 65, which was considered a haploid (Figure 1f). The genome size for this haploid okra was estimated at 1.45Mbp. This plant was feeble, lagging in growth and unfortunately died precociously, but encouraged us to seek for haploid offspring in later samples of reared young okra plants. Such natural haploids, of which a dwarf form of cotton (*Gossypium*) was discovered in 1920 as the first haploid angiosperm with half the normal chromosome complement (Dunwell, 2010), are assumed to result from asexual egg cell (gynogenetic) reproduction (Noumova, 2008). We took advantage of the fact that the diploid hybrid cultivar has a green recessive petiole female parent and a red dominant petiole male (Figure S1) (Portemer *et al*., 2015). Offspring with the green petiole trait lacks the dominant paternal allele and hence can be used as a diagnostic marker for identifying haploid offspring. Accordingly, we selected additional haploid offspring of which one plant was used for sequencing.

**Figure 1.**
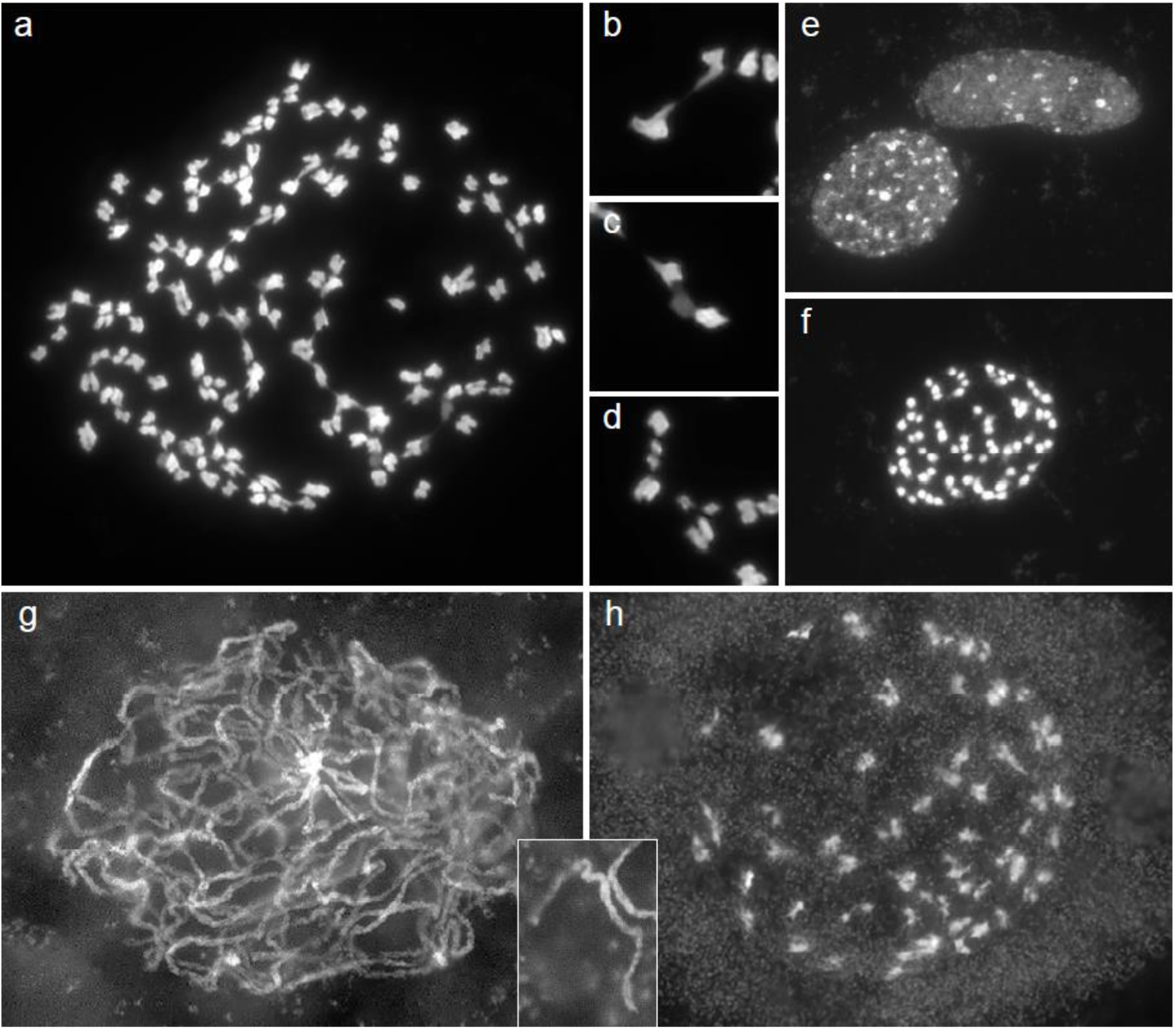
Mitotic cells in root tip meristems. (a) Example of a well spread metaphase complement of a diploid okra plant with 2n=130. (b) Magnification of two chromosomes that are joined by telomere connectives. (c) Chromosome pair with a less fluorescing region, likely representing a decondensed Nucleolar Organizer Region (NOR). (d) chromosomes with small satellites. (e) Two interphase nuclei displaying a striking difference in number of highly condensed chromocenters, regions of the pericentromere heterochromatin and NORs. The top nucleus has about 10 chromocenters; some other nuclei can have more than 100 of such condensed regions. (f) Metaphase complement of a haploid okra plant (2n=65). (g,h) Meiotic chromosomes in pollen mother cells of a diploid okra plant. (g) Cell at pachytene stage. Most of the chromosomes are fully and regularly paired without clear indications for multiple synapsis, pairing loops or pairing partner switches. The brightly fluorescing regions are the pericentromeres, see also the inset between the figures g and h. (h) Cell at diakinesis. A greater part of the chromosomes clearly forms bivalents. Magnification bars in the figures equals 5 µm.

For the analysis of homologous pairing, chiasma formation and chromosome segregation in diploid okra plants we studied male meiosis in pollen mother cells from young anthers (Kantama *et al*., 2017). Pollen mother cells at meiotic stages are filled with fluorescing granular particles in the cytoplasm, which makes it notoriously difficult to see fine details in chromosome morphology. By long enzymatic digestion and acetic acid maturation we still could make the following details visible: pachytene was strikingly diploid-like with clear bivalents showing denser pericentromere regions and weaker fluorescing euchromatin distal parts (Figure 1f). We did not observe clear inversion loops indicative for inversion heterozygosity or pairing partner switches that demonstrate homoeologous multivalents or heterozygous translocation complexes, however the occurrence for such chromosome structure variants could not be excluded. Cell complements at diakinesis displayed that most (if not all) chromosome configurations were bivalents, supporting a diploid like meiosis (Figure 1g). We did not see univalents or laggards at later stages, and pollen were strikingly uniform and well stained (data not shown).

### Okra haploid genome reconstruction

Based on public reports (Benchasri, 2012; Salameh, 2014) and cytological analysis presented above, we applied several technologies for genome reconstruction of the okra haploid individual. 10X Genomics linked read information was used to obtain sequence information in the 100kb to 150kb range. This microfluidics-based technology combines barcoded short-read Illumina sequencing, allowing a set of 150 bp paired-end reads to be assigned to large insert molecules. We produced 800 Gbp of linked read sequencing data from three libraries with an average GC-content of 34% (Table S2). Furthermore, we applied PacBio Circular Consensus Sequencing (CCS), generating 1,400 Gbp of polymerase reads of up to 150kb from circularized insert molecules from three sequence libraries with fragment insert sizes of 10, 14 and 18 Kbp respectively. Polymerase reads were subsequently processed into consensus or so-called HiFi reads with an average sub-read length of approximately 13.6 kb and a sequence error rate less than 1% (Table S2). Over 93% of 1,000 randomly sampled CCS reads had a best BlastN hit to species from the *Malvaceae* family with *Gossypium* ranking first in number of hits, indicating the consistent taxonomic origin, in contrast to the *Abelmoschus* species that are apparently less represented in the NCBI sequence database (Table S3). Furthermore, the organellar DNA content was sufficiently low (Table S4), illustrating the efficiency of our nuclear DNA sample preparations. Upon assembling the HiFi reads with the Hifiasm assembler (Cheng *et al*., 2021), we obtained 3,051 high quality primary contigs with an N50 contig length of 18.9 Mb (Table S5). The incremental sequence assembly displayed in the A50 plot (Figure S2) shows a plateau genome size of approximately 1.35 Gbp, which agrees with the nuclear 2C DNA content. The assembly also resulted in 972 alternative contigs, although their total size of 31 Mbp was small, indicating a highly consistent primary assembly. We nevertheless assessed the origin of alternative contigs, using a taxon annotated GC (TAGC) screen (Kumar *et al*., 2013), providing a means to discriminate between on-target and off-target genomic sequence based on the combined GC content and read coverage and corresponding best matching sequence in annotated databases. The distribution and specific classes of Blast hits indicated that approximately 30% of the alternative contigs could be mapped against annotated sequences (Figure S3), while two thirds were of unknown origin. Alternative contigs had a GC content of 47.6%, which was proportionally higher compared to primary contigs. Furthermore, BlastN hits pointed to a fungal and, to a lesser extent, a bacterial origin. Thus, the smaller sized alternative contigs represented yet a minor contamination in the gDNA sample.

We next physically mapped the genome with BioNano Genomics technology to determine the genome structural organization. We produced 4.88 Tb of unfiltered genome map data with an N50 molecule size of 90.18 kb (Table S2) and a label density of 15.9 per 100kb (Table S6) from nuclei preparations of leaf samples. Size filtering for molecules larger than 100kb left approximately 1.2 Tb of genome mapping data with an N50 molecule size of 206.8 kb (Table S6). Next, molecules, having matching label position and distance, were *de novo* assembled into 216 genome maps with an N50 length of 12.98 Mbp and a total length of 1248.8 Mbp, representing an effective coverage of 375X (Table S6). The genome map size was consistent with the genome sequence assembly size of 1.2 Gbp and thus provided high quality ultra-long-range information for further scaffolding. For that, genome maps were aligned with the *in silico* DLE restriction maps from primary sequence contigs and assembled into higher order scaffolds. The alignment required the cutting of 1 optical map and 4 sequence assembly contigs to resolve conflicts between Bionano maps and sequence contigs respectively, indicating a consistent orientation and order between both. The resulting hybrid assembly was substantially less fragmented, yielding 80 scaffolds with an N50 scaffold size of 18.93 Mb and a total length of 1.19 Gbp, of which the largest scaffold sized more than 29 Mbp (Table 1). Additional scaffolding with 10X Genomics linked reads finally yielded 78 scaffolds (Table 1). Approximately 57% of the individual molecules could be mapped back to the hybrid assembly, suggesting a high confidence genome scaffold.

**Table 1.** NGS assembly and hybrid scaffolding statistics. Sequences were assembled using the Hifiasm assembler and scaffold with Bionano Genomics genome maps. Number of scaffolds obtained with 10X Genomics linked reads is indicated between brackets.

| Assembly statistic | Primary ctgs | Alternative ctgs | Hybrid scfds |
| --- | --- | --- | --- |
| **Ctgs/scfds** | 3,051 | 972 | 80 (78) |
| **Total length** | 1,223.6 Mb | 31.0 Mb | 1,194.5 Mb |
| **Median length** | 32.4 kb | 24.5 kb | 16.320 Mb |
| **Max length** | 25.1 Gb | 652 kb | 29.444 Mb |
| **Min length** | 13.3 kb | 10.3 kb | 125 kb |
| **N50 length** | 10.6 Gb | 30.5 kb | 18.929 Mb |
| **N50 index** | 43 | 126 | 27 |
| **N95 length** | 483 kb | 15.5 kb | 7.206 Mb |
| **N95 index** | 168 | 465 | 64 |
| **GC content** | 34.36% | 47.6% | 33.76% |

### BUSCO analysis and topology of orthologs

To assess the completeness of the genome assembly we screened for BUSCO gene presence/absence (Simaõ, 2015) with 2,326 reference orthologs from the eudicots_odb10 dataset. Based on a best tBlastN hit, 2270 (98%) core genes in 78 scaffolds were detected as ‘Complete’ orthologs (Table 2). Of these, 284 (12.2%) genes were detected as a single copy ortholog. A very small amount (0.3%) was classified as ‘Fragmented’, whereas 32 core genes (1.3%) could not be detected, classifying them as ‘Missing’. These missing BUSCO genes were confirmed to be missing in the alternative contigs as well. We further grouped 2004 (86.2%) multiplicated ortholog genes according to their copy number. A majority of 1150 (49.4%) and 843 (36.2%) orthologs were detected as duplicated and triplicated genes respectively. Interestingly, we found seven and three core genes that were quadruplicated (0.7%) and quintuplicated (0.1%) respectively, and detected one septuplicated core gene, pointing to a complex polyploid nature of the okra genome. To get more insight into the sub-genome organization, the genomic position and topology of ortholog gene copies was assessed. This revealed duplicated BUSCO genes predominantly occurring on two contigs, whereas only a single duplicated ortholog was detected on one contig (Table S7). Both tandem copies were spaced within 1 kb, thus likely representing paralogous genes. Out of 800 triplicate BUSCO’s, 794 (99%) occurred on three contigs, representing three alleles of the same core gene, whereas only six sets (1%) of triplicate core genes were positioned on two contigs. Also, quadruplicate, quintuplicate and septuplicate BUSCO’s mainly occurred on three contigs. The ortholog copies of these groups manifested in a tandem configuration, probably also representing paralogs. Tandemly arranged ortholog copies on the same contigs always showed less sequence distance than between ortholog copies on different contigs. Moreover, the ortholog copies of the septuplicate core gene were dispersed over three contigs. Of these, one contig displayed a triplet, whereas the two other contigs each contained gene copies in a doublet configuration. The triplet consisted of two closely related and one more distantly related paralog. The observed distribution of BUSCO orthologs thus pointed to at least two sub-genomes. However, at this point we could not rule out a higher number of sub-genomes, which might not be discriminated because of a low allelic diversity. Considering a sub-genomic organization for okra, we presume that 284 ‘single copy’ BUSCO genes are either truly unique, or they are maintained as gene copies with indistinguishable alleles.

**Table 2.** Detection of ortholog core genes. Genome assemblies at different coverage levels were analysed to assess the assembly completeness. BUSCO classes are shown as single copy or multiplicated ortholog. Multiplicated orthologs are subdivided into additional copy classes as indicated. For each assembly coverage level BUSCO counts in primary, alternative, and all contigs are shown in absolute numbers and percentages of total expected orthologs (between brackets), or n.d (not determined).

| Class | BUSCO statistics |  |  |  |  |
| --- | --- | --- | --- | --- | --- |
| **Assembly version** | 1 | 2 | 3 | 4 | 4 |
| **Coverage** | 20X | 84X | 95X | 95X | 95X |
| **Ctgs/scfds** | all | all | all | primary | alternative |
| **Complete** | 2,288 (98.3%) | 2,271 (97.7%) | 2,269 (97.6%) | 2,288 (98.4%) | 66 (2.8%) |
| **Single copy** | 266 (11.4%) | 311 (13.4%) | 313 (13.5%) | 284 (12.2%) | 66 (2.8%) |
| **Multiplicated** | 2,022 (86.9%) | 1,960 (84.3%) | 1,956 (84.1%) | 2,004 (86.2%) | 0 (0%) |
| **Duplicated** | n.d | n.d | n.d | 1,150 (49.4%) | 0 (0%) |
| **Triuplicated** | n.d | n.d | n.d | 843 (36.2%) | 0 (0%) |
| **Quadruplicated** | n.d | n.d | n.d | 7 (0.3%) | 0 (0%) |
| **Quintuplicated** | n.d | n.d | n.d | 3 (0.1%) | 0 (0%) |
| **Sextuplicated** | n.d | n.d | n.d | 0 (0%) | 0 (0%) |
| **Septuplicated** | n.d | n.d | n.d | 1 (0.04%) | 0 (0%) |
| **Fragmented** | 5 (0.2%) | 5 (0.2%) | 6 (0.3%) | 6 (0.3%) | 9 (0.4%) |
| **Missing** | 33 (1.5%) | 50 (2.1%) | 51 (2.1%) | 32 (1.3%) | 2,251 (96.8%) |
| **Total** | 2,326 | 2,326 | 2,326 | 2,326 | 2,326 |

Several examples of BUSCO duplication levels in homozygous and heterozygous diploids as well as in auto and alloploids have previously been presented for other species. For example in allotetraploid (2n=4x=38) *Brassica napus* 90% of BUSCO’s are duplicated, whereas only 14.7% in its diploid relative *Brassica campestris* (2n=2x=18) or Chinese cabbage is duplicated (Table S8). The allotetraploid white clover (2n=4x-32) (*Trifolium repens*), that has suggested to be evolved from 2 related diploid species *T. occidentale* (2n=2x=16) and *T. pallescens* (2n=2x=16), has 57% of duplicated BUSCO’s, compared to 10% and 11% of BUSCO duplicates in its diploid ancestral relatives respectively (Griffiths *et al*., 2019). Duplicated BUSCO’s in hexaploid bamboo *B. amplixicaulis* (2n=6x=72) has increased to 57% compared to 35% in its diploid bamboo relative *O. latifolia* (2n=2x=22) (Guo *et al*., 2019). Significant differences were also observed in BUSCO scores between heterozygous and homozygous diploid *Solanaceae*. For example the heterozygous *S. tuberosum* RH potato (2n=2x=24) showed 74.1% of its BUSCO’s duplicated, which was significantly more than 4.3% of duplicated BUSCO’s detected in diploid inbred *S. chacoense* M6 potato (2n=2x=24), and 9.5% detected in autotetraploid inbred *S. tuberosum* potato (Kyriakidou *et al*., 2020). Considering these trends, the BUSCO copy numbers in the okra genome likely point to an allopolyploid nature. Our results confirm the allopolyploid nature of okra as reported by Joshi and Hardas (1956).

### K-mer counts and smudgeplot analysis

To estimate heterozygosity level, repetitiveness, genome size and ploidy levels, we determined kmer counts from raw Illumina and HiFi reads. We compared the 21-mers counts for okra to two related allotetraploid cotton species (*G. barbadese* and *G. hirsutum*), each having a genome of approximately 2.3Gb, and subsequently visualized the readout with Smudgeplot (Ranallo-Benavidez *et al*., 2020) (Figure S4). The k-mer based genome size estimation for the haploid okra amounted to 1.2 Gbp, approximating the NGS assembly size. Approximately 75% of the okra kmers was assigned to an ‘AB’ type (Figure S4). Thus, a major part of the okra genome apparently behaved as a diploid, which is consistent with our cytological observations of a diploid like meiosis, and also coincides with the high number of duplicated BUSCO scores. Approximately 15% of all kmer pairs showed a triploid behaviour (‘AAB-type’). Furthermore, the ‘AAAB’-kmer type seemed more prominent than the ‘AABB’-kmer type. Previously, published kmer readouts for *G. barbadense* and *G. hirsutum* showed that at least 50% of the cotton genomes behaved like a diploid, almost a quarter displayed a triploid behaviour and 14% of the kmers showed tetraploid characteristics. Furthermore, cotton kmer distributions showed the ‘AAAB’ type more frequently occurring than the ‘AABB’ kmer type, which was suggested to be a characteristic for alloploids (Ranallo-Benavidez *et al*., 2020). Thus Genomescope and Smudgeplot readouts for okra point to an allopolyploid nature of the genome, though less complex and smaller sized than anticipated.

### Transcriptome profiling and structural annotation

In addition to sequencing the nuclear genome, we generated approximately 1.2 Tbp of IsoSeq data from multiple tissues including leaf, flower buds and immature fruits to profile the okra transcriptome. The polymerase mean read lengths of up to 86kb benefitted the processing into high quality CCS reads with a mean length of 4.7 kb (σ=378 bp), indicating the efficient full length transcript sequencing (Table S8). The CCS reads were used as transcript evidence for okra gene modelling with the Augustus and GeneMark algorithms from Braker2. We subsequently annotated the 78 largest okra scaffolds with 130,324 genes. Predicted genes had an average length of 2537 nts, whereas the average per gene intron and coding sequence lengths amounted to 307 nts and 2497 nts respectively (Table 3). Coding regions showed low sequence diversity, as only 1109 and 8127 SNPs could be called from full-length transcripts of the okra haploid and an unrelated diploid individual respectively (Table S9). Strikingly, the discovered genes appeared to be predominantly located at the distal ends of scaffolds, gradually decreasing in abundance toward more centrally positioned scaffold domains. In contrast, LTR retrotransposons were more abundant in centrally located scaffold domains, while less frequently represented in the distal ends (Figure 2). A comparable distribution of gene and LTR-retrotransposon regions has been observed for other species such as tomato. The gene and LTR-retrotransposon predicts a heterochromatin organization of pericentromere heterochromatin and distal euchromatin as shown in Figure 1g and inset. This pattern is common in species with small or moderate chromosome size like Arabidopsis, rice and tomato. Apparently, okra also has relatively small sized chromosomes as is substantiated by our cytological observations. The gene-rich regions predominantly occur in euchromatin rich distal chromosome ends and gradually decrease towards the repeat rich more condensed pericentromeric heterochromatin, whereas LTR-retrotransposons were more frequently distributed in pericentromeric heterochromatin (Peters *et al*., 2009;Tomato Sequencing Consortium, 2012; Aflitos *et al*., 2014). Our observations thus suggest a similar chromatin architecture for okra chromosomes. Approximately 51% of the assembled genome was found in the repetitive fraction with 20.26% of repeats unclassified (Table 3). A substantial part (28.69%) consisted of retroelements, of which 24.8% and 1.17% was identified as retrotransposon and DNA transposon respectively. *Gypsy* and *Ty1*/*Copia* retroelements, spanning 15.26% and 9.02% of the assembled genome respectively, appeared to be most abundant (Table 3).

**Figure 2.**
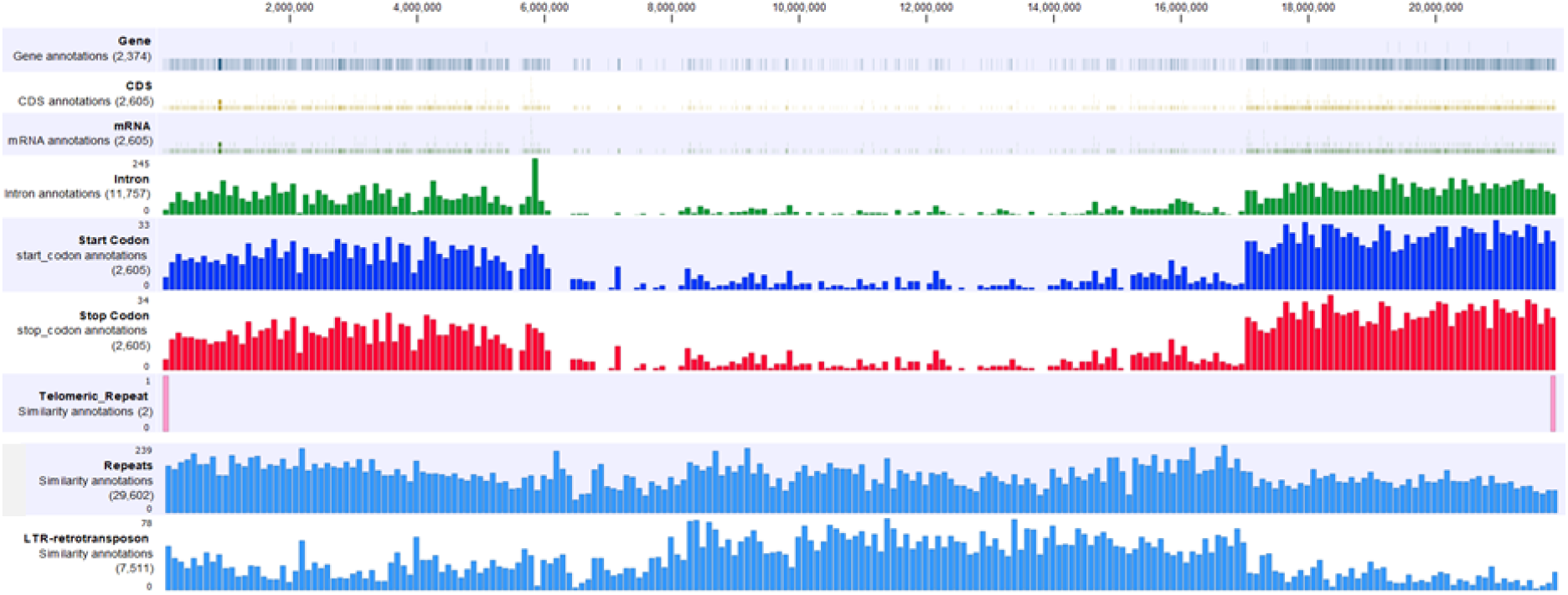
Structural annotation. Annotation feature classes for a 25 Mbp okra scaffold and coordinate positions are indicated at the left and top side of the plot respectively. Bar heights in each row corresponds to the relative frequency of genic, non-genic, and repeat class per scaffold segment of approximately 115 kb.

**Table 3.**
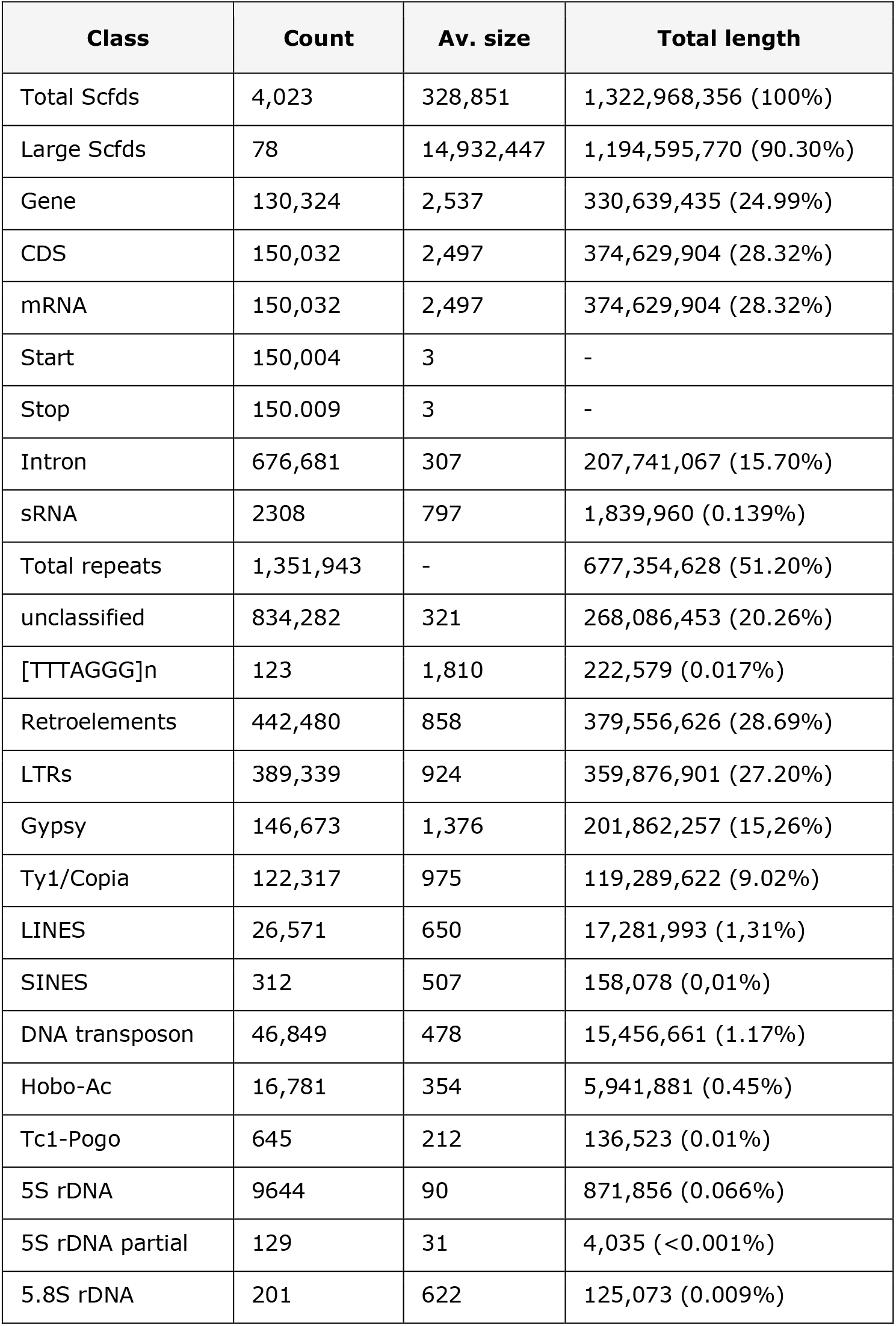
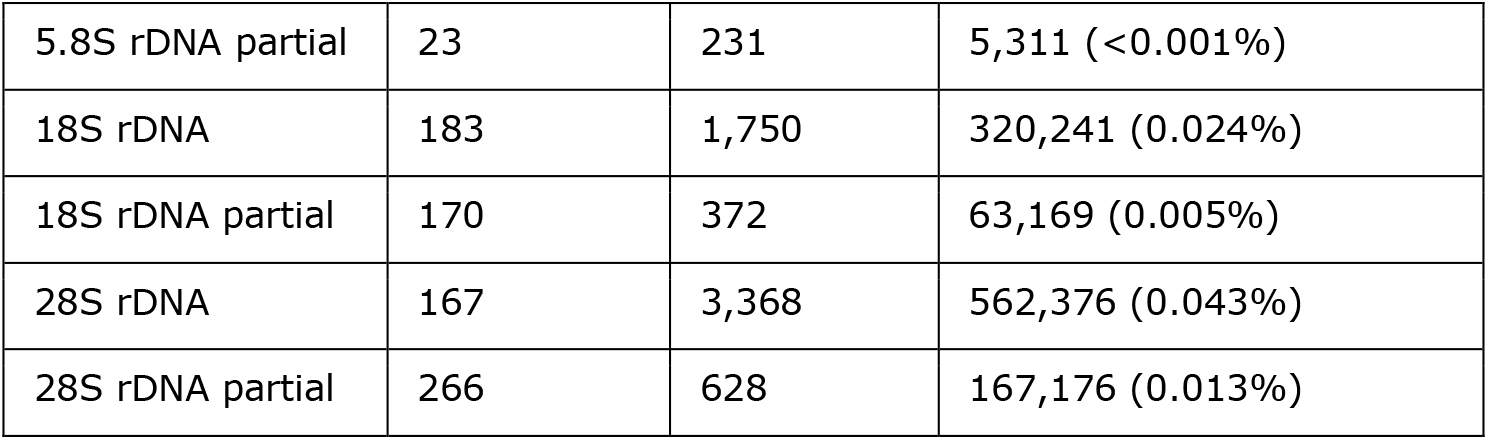
Structural annotation for the 78 largest okra scaffolds. Features are classified into genic and repeat elements as indicated. Statistics are in nucleotide length and in fractions of total scaffold length.

| **Class** | **Count** | **Av. size** | **Total length** |
| --- | --- | --- | --- |
| Total Scfcs | 4,023 | 328,851 | 1,322,968,356 (100%) |
| Large Scfcs | 78 | 14,932,447 | 1,194,595,770 (90.30%) |
| Gene | 130,324 | 2,537 | 330,639,435 (24.99%) |
| CDS | 150,032 | 2,497 | 374,629,904 (28.32%) |
| mRNA | 150,032 | 2,497 | 374,629,904 (28.32%) |
| Start | 150,004 | 3 | - |
| Stop | 150,009 | 3 | - |
| Intron | 676,681 | 307 | 207,741,067 (15.70%) |
| sRNA | 2308 | 797 | 1,839,960 (0.139%) |
| Total repeats | 1,351,943 | - | 677,354,628 (51.20%) |
| unclassified | 834,282 | 321 | 268,086,453 (20.26%) |
| [TTTAGGG]n | 123 | 1,810 | 222,579 (0.017%) |
| Retroelements | 442,480 | 858 | 379,556,626 (28.69%) |
| LTRs | 389,339 | 924 | 359,876,901 (27.20%) |
| Gypsy | 146,673 | 1,376 | 201,862,257 (15.26%) |
| Ty1/Copia | 122,317 | 975 | 119,289,622 (9.02%) |
| LINES | 26,571 | 650 | 17,281,993 (1.31%) |
| SINES | 312 | 507 | 158,078 (0.01%) |
| DNA transposon | 46,849 | 478 | 15,456,661 (1.17%) |
| Hobo-Ac | 16,781 | 354 | 5,941,881 (0.45%) |
| Tc1-Pogo | 645 | 212 | 136,523 (0.01%) |
| 5S rDNA | 9644 | 90 | 871,856 (0.066%) |
| 5S rDNA partial | 129 | 31 | 4,035 (<0.001%) |
| 5.8S rDNA | 201 | 622 | 125,073 (0.009%) |
| 5.8S rDNA partial | 23 | 231 | 5,311 (<0.001%) |
| 18S rDNA | 183 | 1,750 | 320,241 (0.024%) |
| 18S rDNA partial | 170 | 372 | 63,169 (0.005%) |
| 28S rDNA | 167 | 3,368 | 562,376 (0.043%) |
| 28S rDNA partial | 266 | 628 | 167,176 (0.013%) |

The repeat screening also revealed stretches of the Arabidopsis telomere TTTAGGG motif, flanking gene rich regions at distal scaffold ends (Figure 3). In plants such repeats usually occur in high copy numbers at the distal ends of chromosomes, constituting telomeres that protect the terminal chromosomal DNA regions from progressive degradation and preventing the cellular DNA repair mechanism from mistaking the ends of chromosomes for a double stranded break. Indeed, we found blocks of TTTAGGG units in high copy numbers positioned at both ends for 49 scaffolds, whereas 25 scaffolds had a telomere repeat block at one end, in total 123 telomeres at the end of 130 chromosome arms. This repeat distribution suggested full length chromosome scaffolds and capturing the majority of 65 chromosome ends of the haploid okra genome and again confirms the relatively small sized okra chromosomes. Besides long distal blocks we also detected short interstitial TTTAGGG blocks. These interspersed non-telomeric short TTTAGGG repeats possibly reflect footprints of internalized telomeres that may have arisen from end-to-end fusion of chromosomes (Baird, 2017), possibly representing hallmarks of chromosomal speciation upon allopolyploidization of okra. Another substantial fraction of repeats originated from ribosomal genes. BlastN analysis, using *Gossypium hirsutum* ribosomal gene query sequences, clearly showed an 18S, 5.8S and 28S rRNA gene block arrangement in okra. Two clusters are located at scaffolds ends, though not coinciding with, or flanking telomere blocks. Another two clusters are positioned toward the scaffold centre, and three scaffolds almost entirely consist of 18S-5.8S-28S gene clusters. These scaffolds do not contain 5S rRNA clusters. Instead, 5S rRNA genes are organized in clusters separated from the 18S-5.8S-28S units, clearly indicating an S-type rRNA gene arrangement (Goffová and Fajkus, 2021). In total we found four 5S rRNA clusters on four different scaffolds of which the largest cluster consisted of almost 8,700 copies tandemly arranged on a single scaffold (Table S10). Signatures of underlying chromosome evolution involving telomere fusion at ribosomal gene clusters were not apparent though, as we did not encounter interstitial telomere repeats in rRNA clusters.*Genetic diversity in okra*

**Figure 3.**
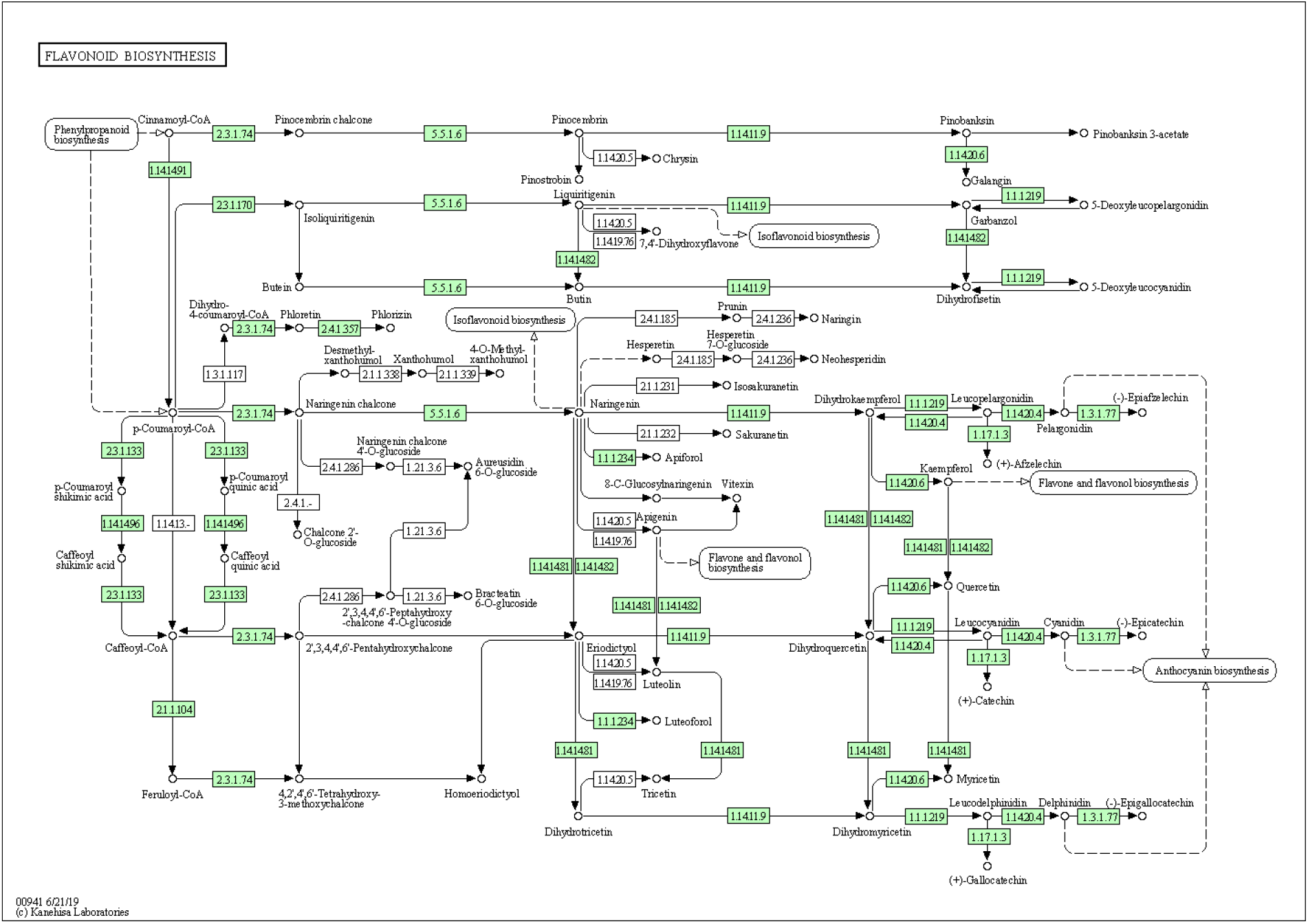
The flavonoid KEGG bio-synthesis pathway in *Abelmoschus esculentus*. Putative okra enzyme coding genes for which a bi-directional best hit was found to reference pathway enzymes are shown with coloured EC identifiers.

To assess the genetic diversity in public okra germplasm, we used the okra reference genome to call single nucleotide polymorphisms (SNPs) from several publicly availaparamount importance to understand plant metabble Illumina RNA-seq datasets that we retrieved from the short-read archive (SRA) (https://www.ncbi.nlm.nih.gov/sra). Most of these samples represent accessions originating from the Indian and Chinese parts of the Asian continent. The average map rate for a panel of 11 samples was 93.2% ± 5.6% (Data S1). The combined samples cover 20-25% of the reference genome, which is in line with the 26.6% genic portion of the genome and suggesting a faithful structural annotation of the reference genome. The unusual high coverage for the ‘Xianzhi’ dataset (∼65%) possibly was due to a deviating library preparation or DNA contamination, while the lower coverage for the ‘IIHR-299’, ‘Mahyco Arka Abhay’, and ‘Commercial’ samples (10-12%) was likely due to their smaller data size. The total panel size comprised 412,185 loci, for which 690,145 SNPs were detected from coding regions of okra genes. The “Commercial” sample yielded only 1,741 unique SNPs with reads mapping to 12.5% coverage of the reference genome. This SNP rate is substantially lower compared to “Mahyco Arka Abhay” and “IIHR-299”, while these 3 samples cover approximately equal portions of the genome, suggesting that the ‘Commercial’ breeding line apparently shares a large part of its ancestry with the reference cultivar. However, difference in tissue types, growth conditions, data generation and processing workflows complicate direct sample comparison. Nevertheless, most of the samples attain 20,000 unique SNPs and only ‘Arka Anamika’ and ‘Danzhi’ exceed this level. Although we could not yet assess the genetic diversity in the non-genic portion, the okra accessions in the panel apparently represent a low genetic diversity.

### Allopolyploid composition of okra

We attempted to divide okra subgenomes by searching for patterns based on hierarchical clustering of repeat k-mer counts without a supposition of ancestral species. We compared the clustering results for okra with two k-mer test sets, of which the first comprised an artificially constructed hybrid genome, consisting of merged tomato (*S. lycopersicum* cv. Heinz 1706) and a diploid potato (*S. tuberosum* cv. Solyntus) genomes. The second set was generated from the allotetraploid cotton genome (*Gossypium hirsutum*). As expected the cluster map for the artificial hybrid dataset separated into two distinct subclusters, each representing repetitive k-mers from 12 tomato and potato chromosomes (Figure S5). In addition, the cotton genome also clustered into two groups, clearly separating the repetitive kmers generated from the subgenome A and D chromosomes (Hu *et al*., 2019). Applying the same method to the okra reference genome, of which five smallest scaffolds were removed, yielded two distinct clusters of 30 and 43 scaffolds with a length of 636 Mbp and 557 Mbp, respectively (Figure S6). Scaffolds from cluster 1 are overall larger than from cluster 2 (Figure S7). Apparently the repeat k-mer pattern for okra points to two distinct subgenomes and, together with the apparent absence of multivalent pairing of metaphase chromosomes, suggests an allotetraploid nature of *Abelmoschus esculentus*. In addition, we could not find clear evidence of erosion, as the clusters had comparable BUSCO completeness scores of 90.2% and 89.3%, with duplicate rates of 21.8% and 19.4% respectively. This suggests a relatively recent hybridization of ancestral species, yet without clear evidence for an emerging dominant subgenome.

We subsequently aligned the clustered scaffolds to further investigate the orthology between the two subgenomes. Although we found only partial alignments and several inversions, there is substantial homoeology between scaffolds (figure S8), confirming in general there are no homoeologs within a single cluster. Specifically, the network of scaffolds displayed the portion of BUSCO genes shared between scaffolds (figure S9), indicates the strong homoeology between scaffolds of cluster 1 and 2, corroborating the alignment dot-plot.

Further repeat annotation reveals only 784 out of 22,100 repeat classes are present in >90% of scaffolds in both clusters. A small number of 14 repeat classes are present in more than 90% of the scaffolds in either cluster, while occurring in less than 10% of scaffolds in the other. These include cluster specific repeat classes, occurring in high copy numbers in cluster 1 but not in cluster 2 and vise versa (Figure S10). Although such repeats do not show similarity to known repeats and we can only speculate about their origin, they are clearly related to different ancestral progenitor species, further pointing to an allotetraploid nature of okra.

### Candidate genes assigned to phenylpropanoid, flavonoid, and flavone and flavonol biosynthesis pathways

Polyphenols represent one of the most ubiquitous class of secondary metabolites in okra fruits. An important subclass of polyphenols are flavonoids, of which myricetin, quercitin, isoquercitrin and quercetin-3-O-gentiobioside derivatives have been implicated in antidiabetic activity (Liu *et al*., 2005; Lei *et al*., 2017; Wu *et al*., 2020; Peter *et al*., 2021). Myricitin was previously detected in *Abelmoschus moschatus* (Liu *et al*., 2005). Recently, the bioactive phytochemicals isoquercitrin and quercetin-3-O-gentiobioside, and to a lesser extent also rutin and catechin, were detected as the major phenolic compounds in okra fruits (Wu *et al*., 2020). Their biosynthesis in the flavonoid and, flavone and flavonol biosynthesis pathways (KEGG reference pathways 00941 and 00944) is thought to start with p-coumaroyl-CoA and cinnamoyl-CoA precursors that are synthesized in the phenylpropanoid pathway (KEGG reference pathway 00940). To find putative enzyme coding okra genes that may function in phenylpropanoid, flavonoid, flavone and flavonol biosynthesis, 142,571 extracted amino acid query sequences from predicted okra genes and putative splice variants were mapped against the manually curated KEGG GENES database, using the KEGG Automatic Annotation Server (KAAS) (KAAS - KEGG Automatic Annotation Server (genome.jp) (Moriya *et al*., 2007). We identified 33,641 amino acid sequences that could be assigned to 395 KEGG metabolic pathway maps based on a best bi-directional hit (BBH). Currently, in total there are N=1302 manually annotated from *Malvaceae* species *Theobroma cacao* (cacao), *Gossypium arboretum, Gossypium hirsutum* (cotton), *Gossypium raimondii*, and *Durio zebithinus* (durian) of which K=99 enzyme coding reference genes are known for the phenylpropanoid (K_1_=36), flavonoid (K_2_=30), or flavone and flavonol (K_3_=33) biosynthesis pathway in KEGG. Of the 33,641 okra amino acid queries, n=47 putative okra orthologs were assigned to the KEGG phenylpropanoid biosynthesis (n_1_=16), flavonoid biosynthesis (n_2_=17) and flavone and flavonol biosynthesis (n_3_=14) pathways respectively, adding up to n=41 distinct putative okra enzyme orthologs. We subsequently assessed the mapping probability of okra orthologs to the reference pathways based on the known *Malvaceae* enzymes and the okra BBH. Mapping confidence values p1=1.43e_-12_, p2=1.15e_-12_, and p3=0.0 pointed to a confident assignment of okra orthologs to phenylpropanoid biosynthesis, flavonoid biosynthesis, and flavone and flavonol biosynthesis pathways respectively. Copy numbers for putative genes possibly involved in the conversion p-coumaroyl-CoA and cinnamoyl-CoA precursors varied extensively. Only a single putative gene orthologous to a 5-O-(4-coumaroyl)-D-quinate_3’-monooxygenase (EC:1.14.14.96) from *Durio zebithinus* (XP_022742205) with an amino acid identity of 93.9% was found, whereas 14 putative orthologs to shikimate O-hydroxycinnamoyltransferase (EC2.3.1.133) were detected, with a highest amino acid identity (94.9%) to the ortholog from *Gossypium arboreum* (XP_017607223). The coverage of okra orthologs mapped to these biosynthesis pathways is shown in figures 3 and S11. The alternative metabolic routes, leading to the biosynthesis of quercitin, myricitin, isoquercitrin, and quercetin-3-O-gentiobioside derivatives in these pathways, involve three critical flavonol synthases CYP74A (flavonoid 3’,5’-hydroxylase, EC:1.14.14.81), CYP75B1 flavonoid 3’-monooxygenase (EC:1.14.14.82) and (FLS) dihydroflavonol,2-oxoglutarate:oxygen oxidoreductase (EC:1.14.20.6). Apparently, putative orthologs for CYP74A and CYP75B1 are encoded by a single copy gene in okra, whereas the FLS oxidoreductase catalytic activity, that is thought to catalyse the conversion of several dihydroflavonol intermediates into quercitin and myricitin, appears represented by 5 putative okra homologs, suggesting that the conversion into quercitin, isoquercitrin, rutin and catechin mainly runs via a dihydrokaempferol intermediate.

## Conclusions

We successfully applied multiple DNA sequencing and scaffolding techniques to reconstruct the 65 chromosomes of a haploid okra sibling. With a final result of 49 scaffolds with telomeres at both ends, illustrating chromosome completeness, and 16 scaffolds with at least 1 telomere end, representing partially resolved chromosomes, the assembly contiguity for okra surpasses that of other crop genome assemblies. PacBio high fidelity (HiFi) reads and availability of a haploid sample were key to the high quality of the reconstructed okra genome. The more pronounced diploid behavior of okra, exemplified by the smudgeplot spectrum showing 75% AB dominance, when compared to the k-mer spectrum for allotetraploid cotton with 50% AB dominance, illustrates a decreased complexity that benefitted the genome reconstruction. This diploid-like nature is also in line with our cytogenetic observations on pollen mother cells at pachytene, showing chromosomes strikingly diploid-like with clear bivalents. The okra genome is characterized by a high chromosome count and relatively small chromosomes that do not extend beyond 30 Mbp in length. The total haploid assembly size of ∼1.2 Gbp apparently is consistent with the slightly larger k-mer based haploid genome size estimation. Mapping of telomeric sequences and gene dense sections toward both scaffold ends, together with repeat dense sections mapping toward more centrally located scaffold sections, is in line with the observed chromatin landscape at pachytene for several okra chromosomes that display pericentromeric heterochromatin and distal euchromatin, which have been shown repeat-rich and gene-rich respectively also in many species genomes. The overall repeat content (57%) is lower than described for *Gossypium raimondii* (70.7%), but higher than for *Theobrama cacao* (29.4%) (Novák *et al*., 2020), whereas these genomes are smaller. Our repeat classification remains incomplete though, due to a lack of diversity in annotated repeat libraries. Furthermore, with over 130,000 putative genes, the gene prediction count seems inflated, compared to other species genomes like A*rabidopsis* and rice (*Oryza sativa*), containing around 38,000 and 35,000 coding, non-coding and pseudogenes respectively. Some increase may be attributed to allopolyploidization. On the other hand, a decreased selective pressure may cause many genes to accumulate *de novo* mutations and convert into pseudogenes (Bird *et al*., 2018). Nevertheless, over 88% of the predicted exons was supported by high-quality long read data and predicted proteins mostly returned partial matches to several different databases, substantiating our gene prediction. In this respect, the structural and functional annotation revealed putative enzyme coding genes that we could map to phenylpropanoid, flavonoid, flavone and flavonol metabolic pathways, likely underlying the biosynthesis of an array of secondary metabolites that have been implicated in dietary and therapeutic bioactivity.

Identification of subgenomes from distinct k-mer repeat profiles point to an allotetraploid composition of the genome. The two subgenomes apparently have unequal chromosome counts, though are similar in total sequence length. The BUSCO scores suggest nearly complete subgenomes, likely reflecting a relatively recent hybridization of two progenitor species without clear evidence for subgenome dominance. Substantial numbers of duplicated BUSCO genes in the subgenomes might reflect hybridization events prior to allotertaploidization. In this respect the additional grouping of k-mers within each of the two k-mer repeat clusters is suggestive and intriguing, though need more detailed analysis. Finally, we conclude that the annotated high-quality genome provides a solid basis for further okra breeding application, including diversity and compatibility screening, and marker development.

## Acknowledgements

We wish to thank Hortigenetics Research of East West Seed (S.E. Asia) Ltd., ENZA Zaden Research and Development B.V., Genetwister Technologies B.V., Nunhems Netherlands B.V., Syngenta Seeds B.V., Takii & Company Ltd., HM.Clause, SA., UPL Ltd., Namdhari Seeds Pvt. Ltd., Maharashtra Hybrid Seeds Co. Pvt. Ltd. and Acsen HyVeg Pvt. Ltd. for providing material and support to the okra genome project.

## Supplementary figures

**Figure S1.**
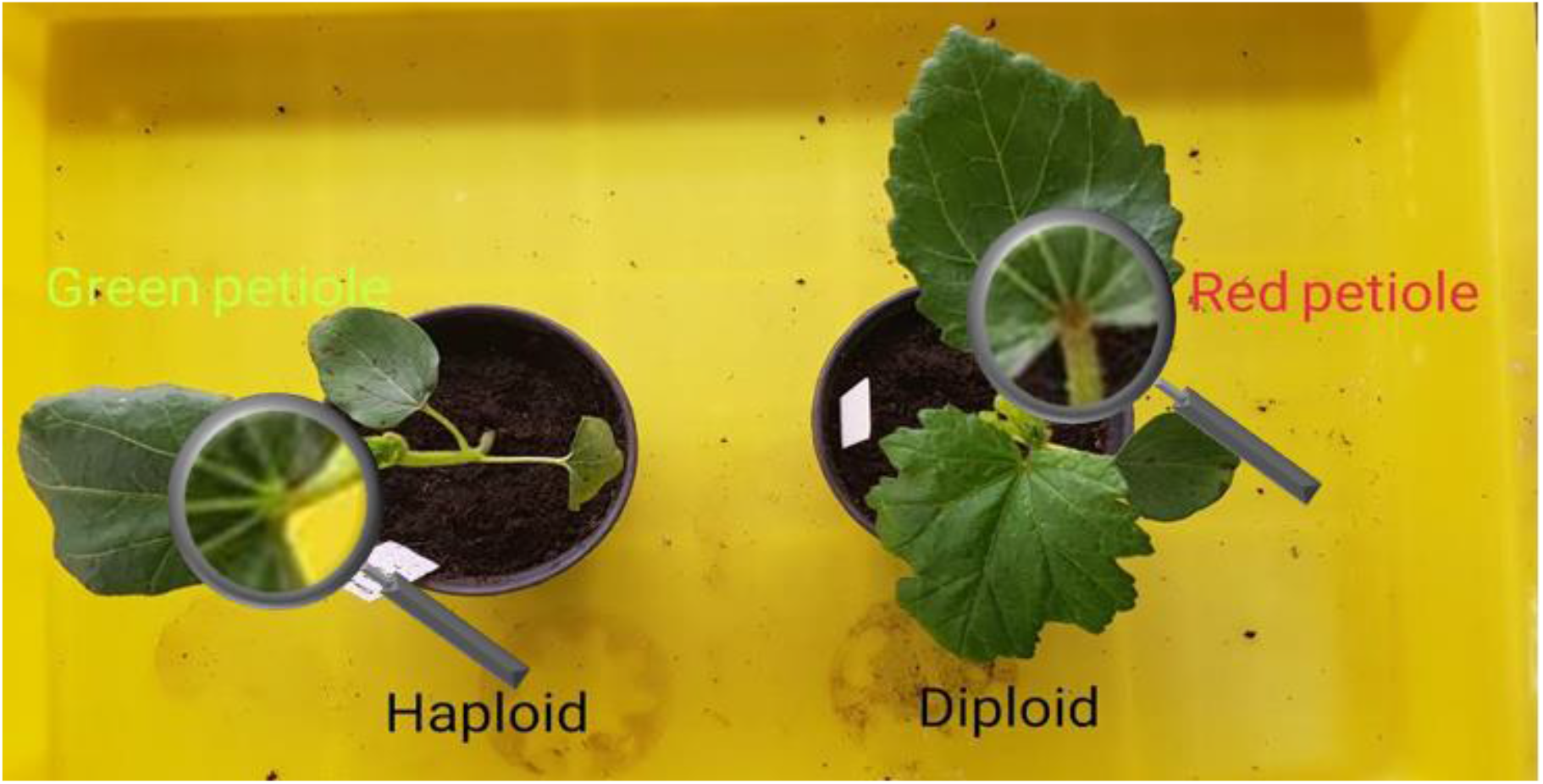
Phenotypes of diploid and haploid Okra plants. The magnifying glass in the image is placed over the position of the green and the red petiole for the haploid (left) and diploid (right) Okra plant respectively.

**Figure S2.**
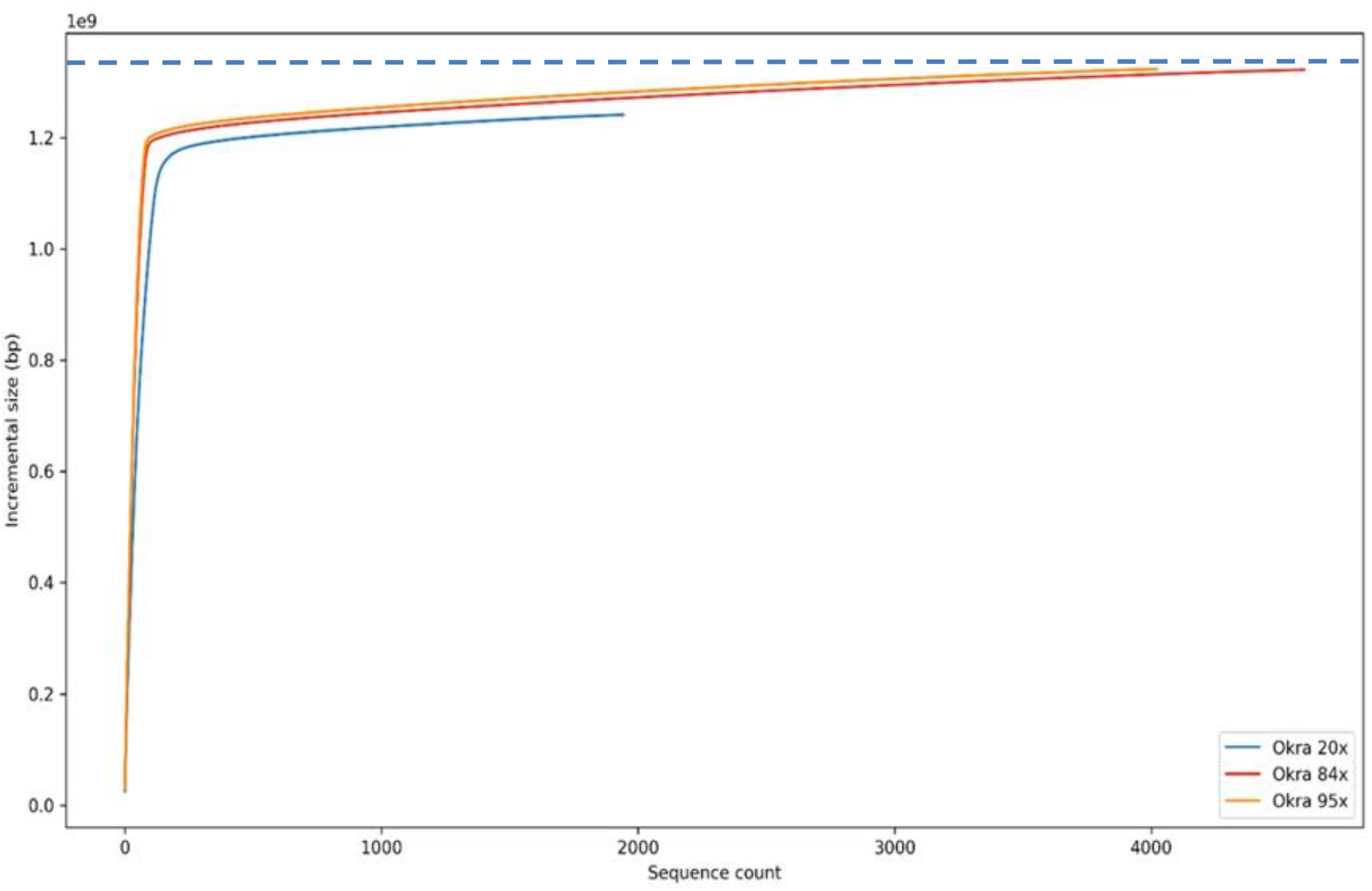
The incremental genome assembly size for Okra. The A50 plot for contigs larger than 100 bp shows the assembly size in Gbp on the y-axis is plotted against the incremental contig count at 20X, 84X and 95X sequence coverage indicated by the light blue, red and orange curve respectively.

**Figure S3.**
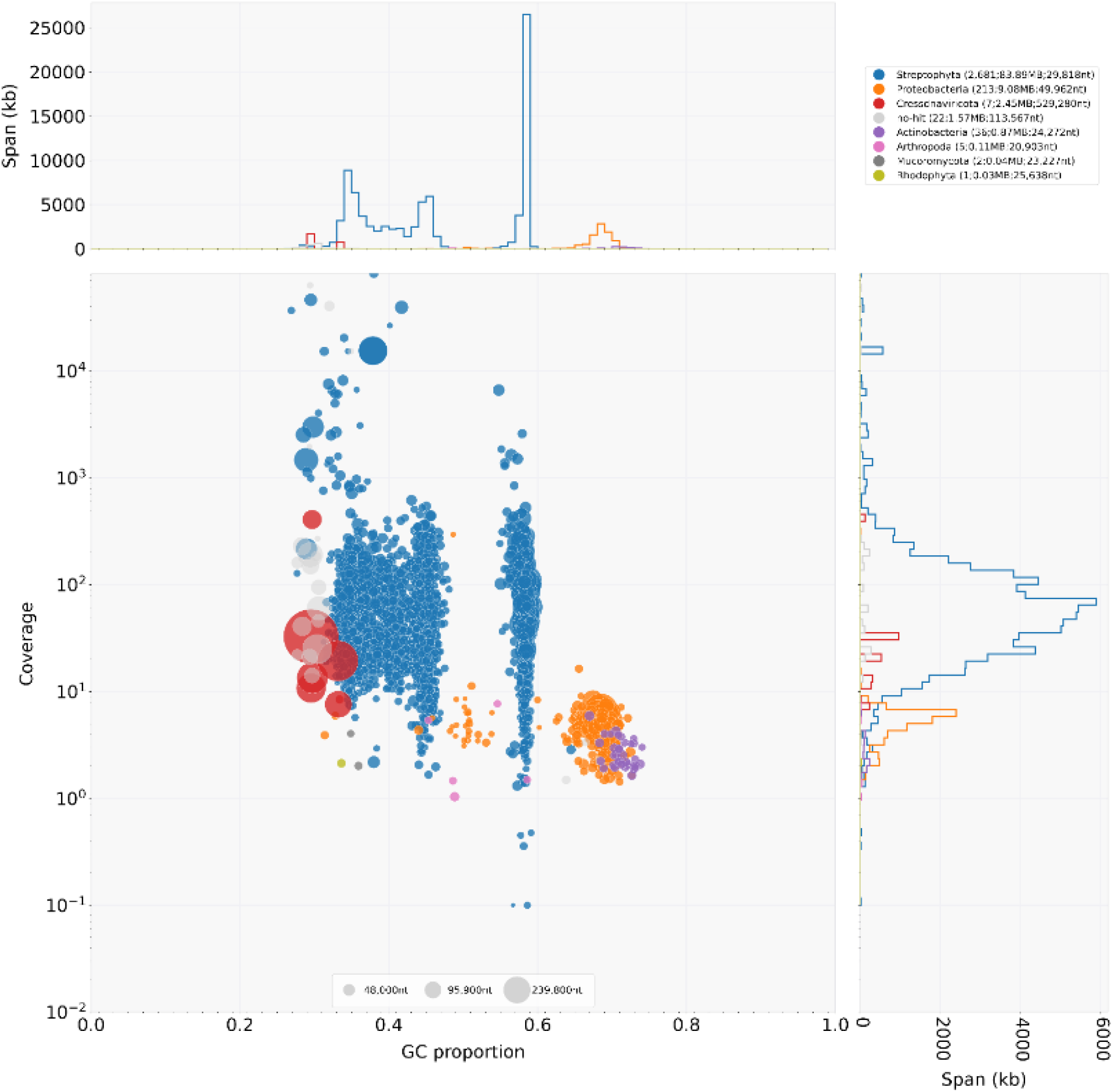
Taxon annotated GC coverage plot. In the low left panel the proportion of GC bases (x-axis) and read coverage (y-axis) for 527 alternative contigs are shown. Each coloured dot in the graph corresponds to a contig. Colours correspond to species classes for which a best BlastN match was found in annotated databases. In the top left graph and the low right graph the relative proportion for each class is depicted with respect to the coverage and GC content, for which colour codes match species classes as indicated in the legend at the top right.

**Figure S4.**
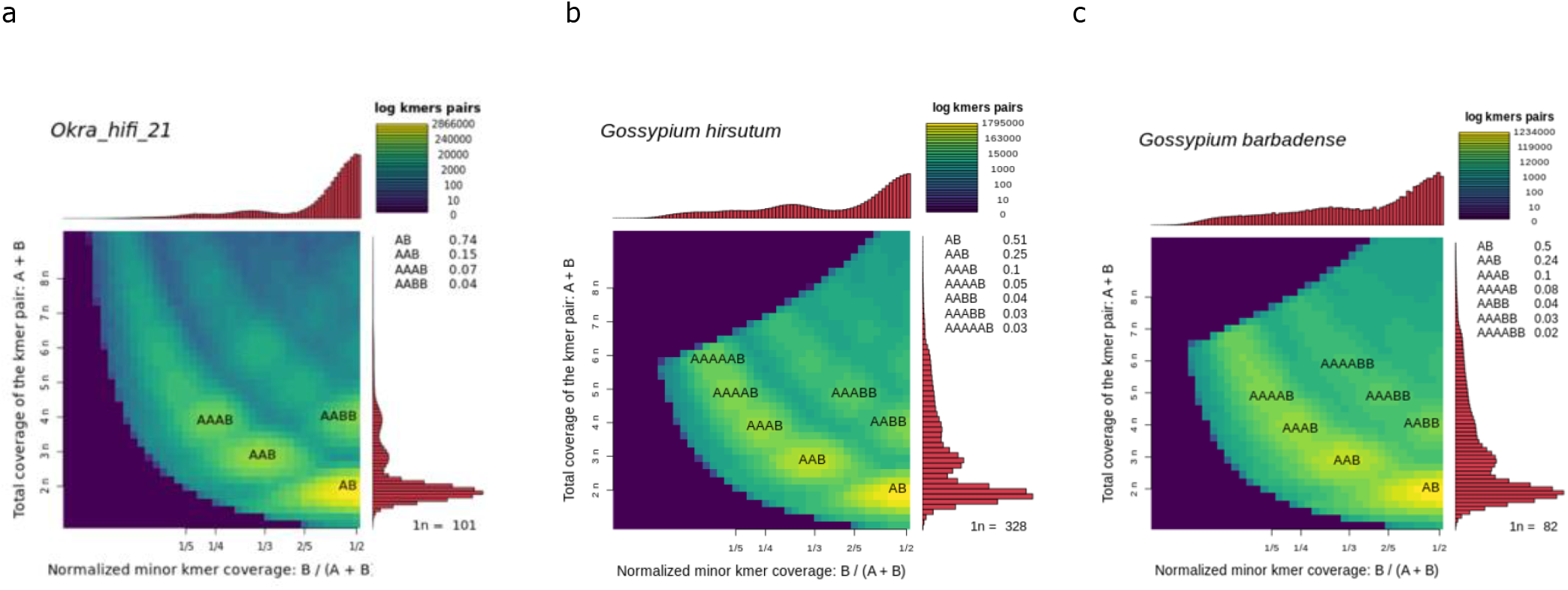
Smudgeplots for and haploid Okra (*Abelmoschus esculentus*) (a), and two allotetraploid cotton species *Gossypium hirsutum* (b), and *Gossypium barbense* (c). Smudgeplots are shown in log scale. The coloration indicating the approximate number of k-mer pairs per bin and the fraction of each kmer type is indicated in the top right legend of each plot.

**Figure S5.**
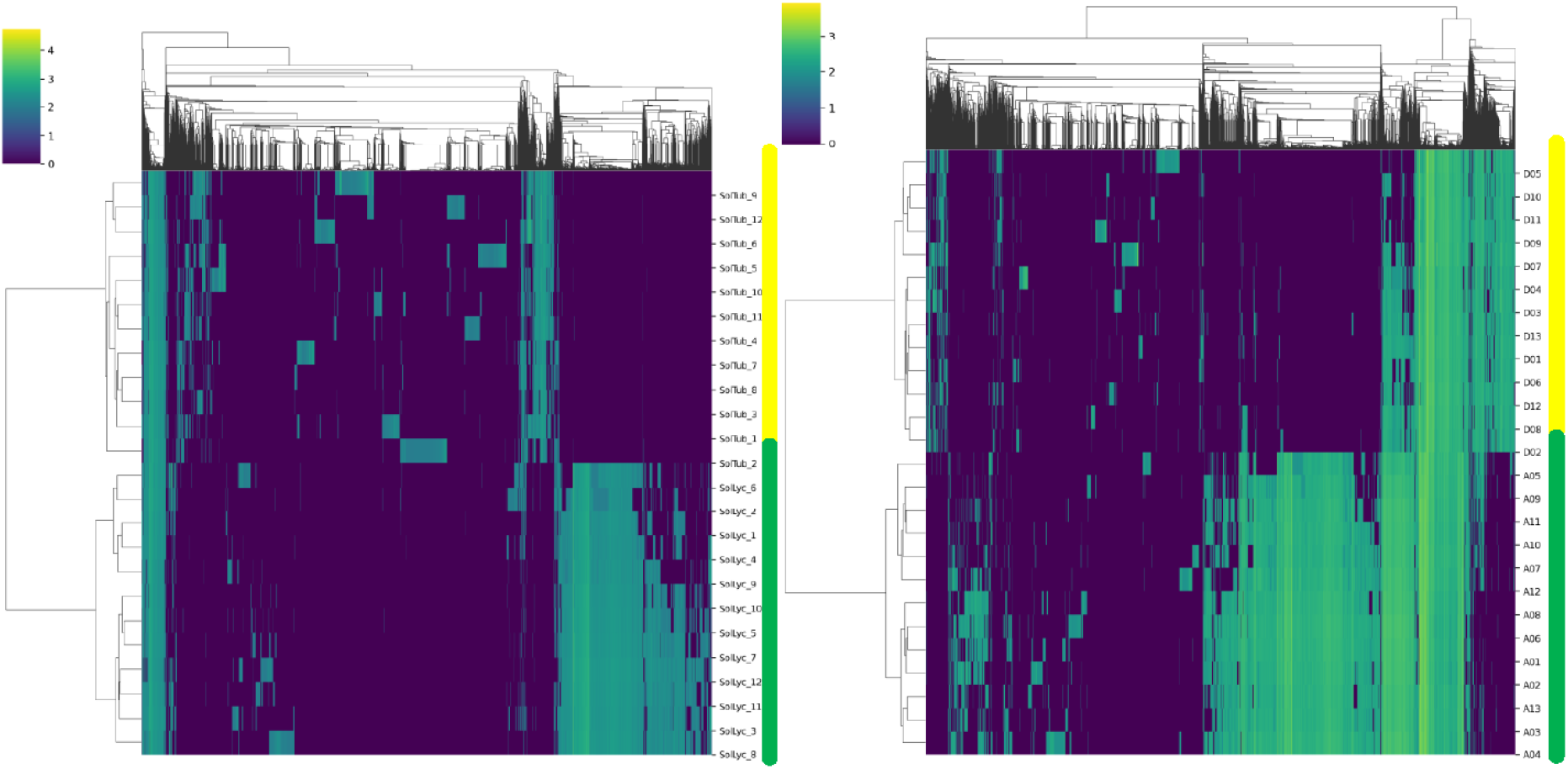
Cluster maps of repetitive 13-mer counts. K-mers generated for an artificial hybrid genome constructed from merged genomes of *S*.*lycopersicum* SL.4.0 and *S. tuberosum* cv. Solyntus (left panel) and from the allotetraploid cotton *G. hirsutum* genome. Yellow and green bars next to the chromosome identifiers mark the potato and tomato chromosomes (left panel) and the cotton chromosomes from the A and D subgenomes (right panel) respectively. Color code bar indicates log10 scaled kmer counts.

**Figure S6.**
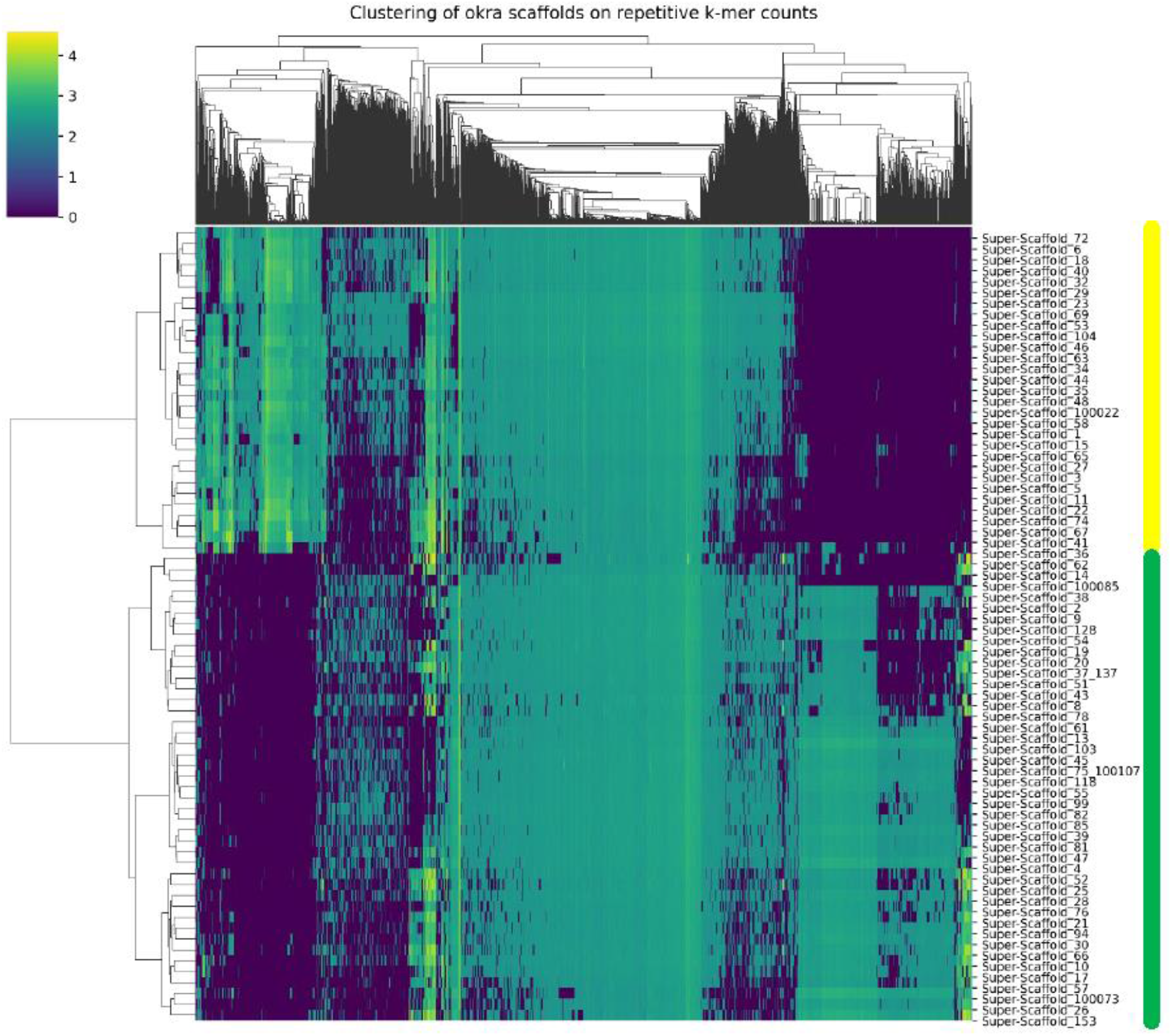
Cluster maps of repetitive 13-mer counts for the okra reference genome. Yellow and green bars next to the identifiers mark the superscaffolds clustering in cluster 1 and 2 respectively. Color code bar indicates log10 scaled kmer counts. A clear separation between cluster 1 and 2 based on repeat count and distinct repeat profile is apparent.

**Figure S7.**
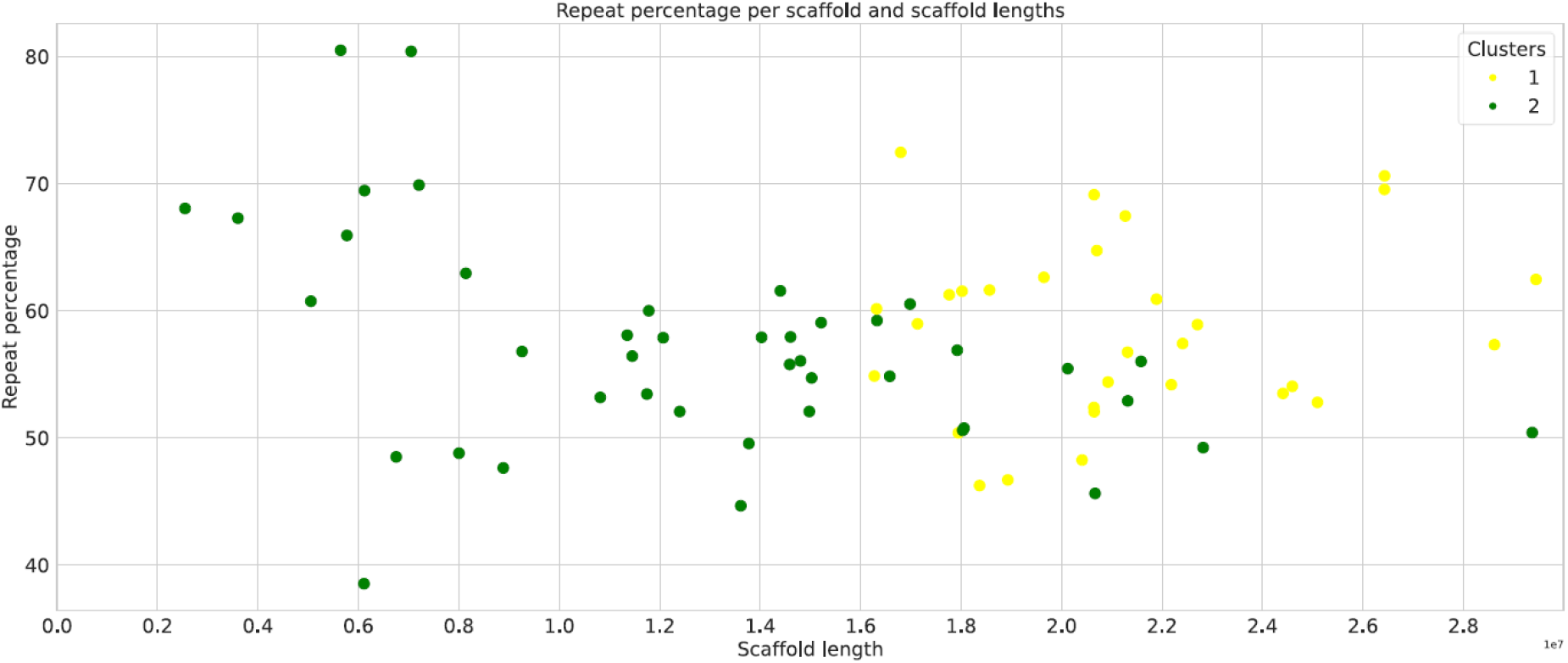
Repeat content analysis in superscaffolds. Superscaffolds assigned to cluster1 and 2 are represented by yellow and green dots respectively, and have been separated by scaffold length (x-axis) and repeat percentage (y-axis).

**Figure S8.**
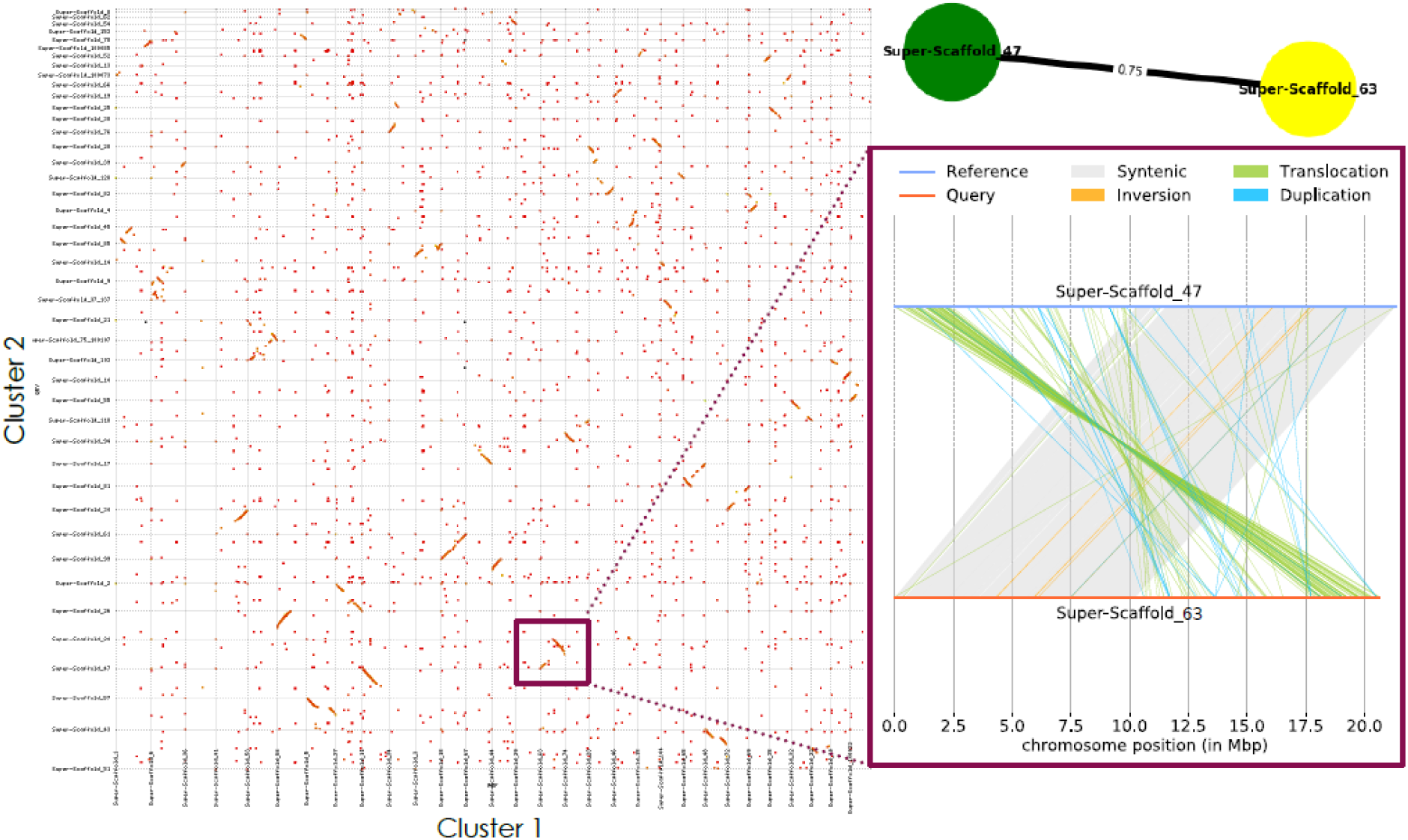
Dot plot alignment of superscaffolds from subgenomes. Superscaffolds are assigned to cluster 1 or 2 according to their kmer clustering profile. The top right graph shows two homoeologous superscaffolds 63 (cluster 1) and 47 (cluster 2) having 75% of BUSCO genes in common. The bottom right alignment detail of the aforementioned superscaffolds are partially syntenic, sharing a large inversion.

**Figure S9.**
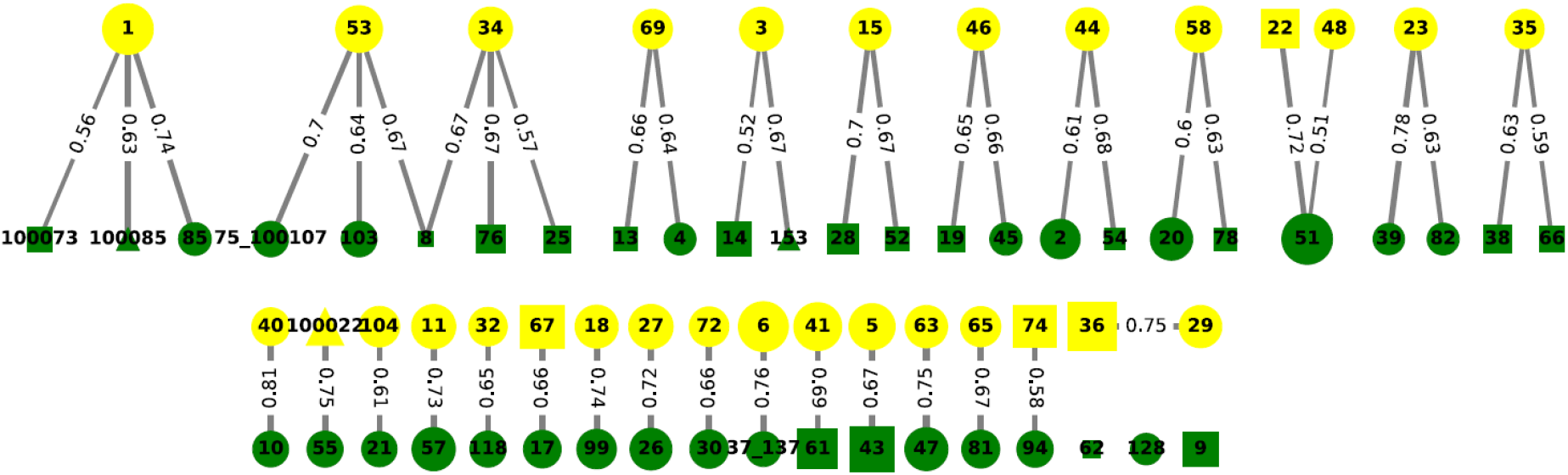
BUSCO connectivity graph. Yellow and green color coded nodes correspond to scaffolds from cluster 1 and 2 respectively. Edges between the nodes indicate the percentage of shared BUSCO genes between each scaffold pair. Note that a single node can have multiple edges. Pairs of scaffolds point to links of homoeology between scaffolds.

**Figure S10.**
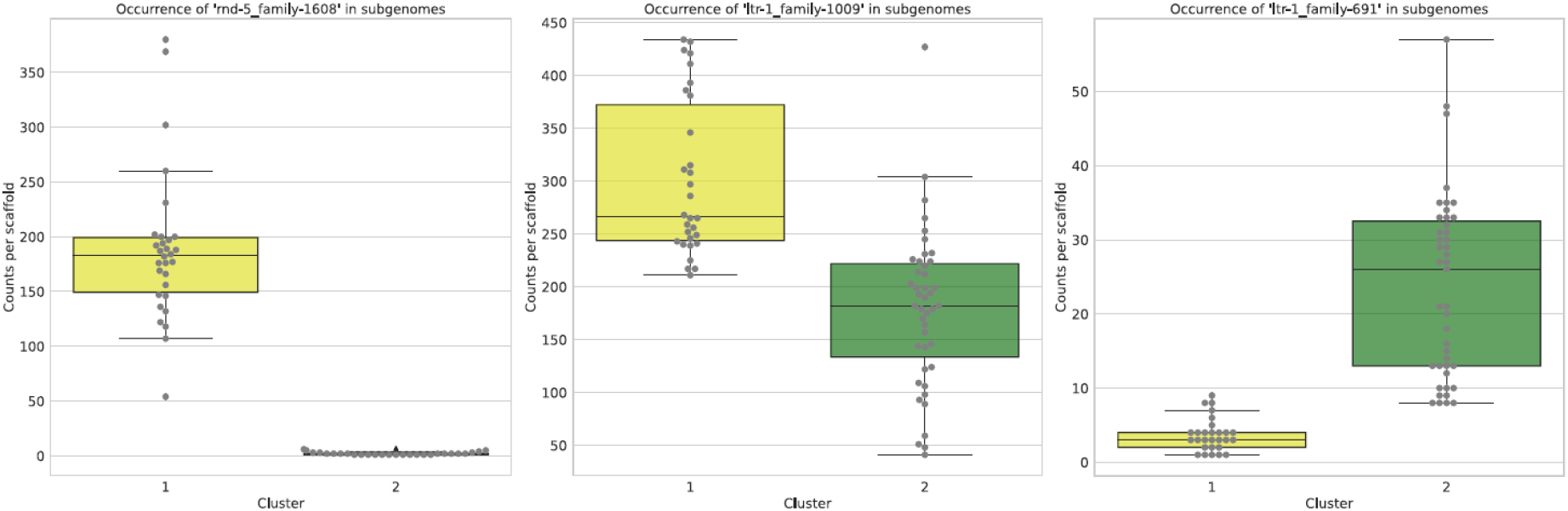
Repeat count of cluster specific and shared repeats between subgenomes. Occurrence of 3 distinct repeat families are shown as counts per scaffold (y-axis) that are divided over distinct clusters 1 and 2 (x-axis). Counts per scaffold are represented by grey dots. The repeat family identifier is indicated above each plot. The left panel shows the occurrence of an unclassified repeat family in cluster 1 specific scaffolds while absent in cluster 2 scaffolds. The right panel shows an unclassified cluster 2 specific repeat family. The unclassified repeat family in the middle graph is cluster unspecific.

**Figure S11.**
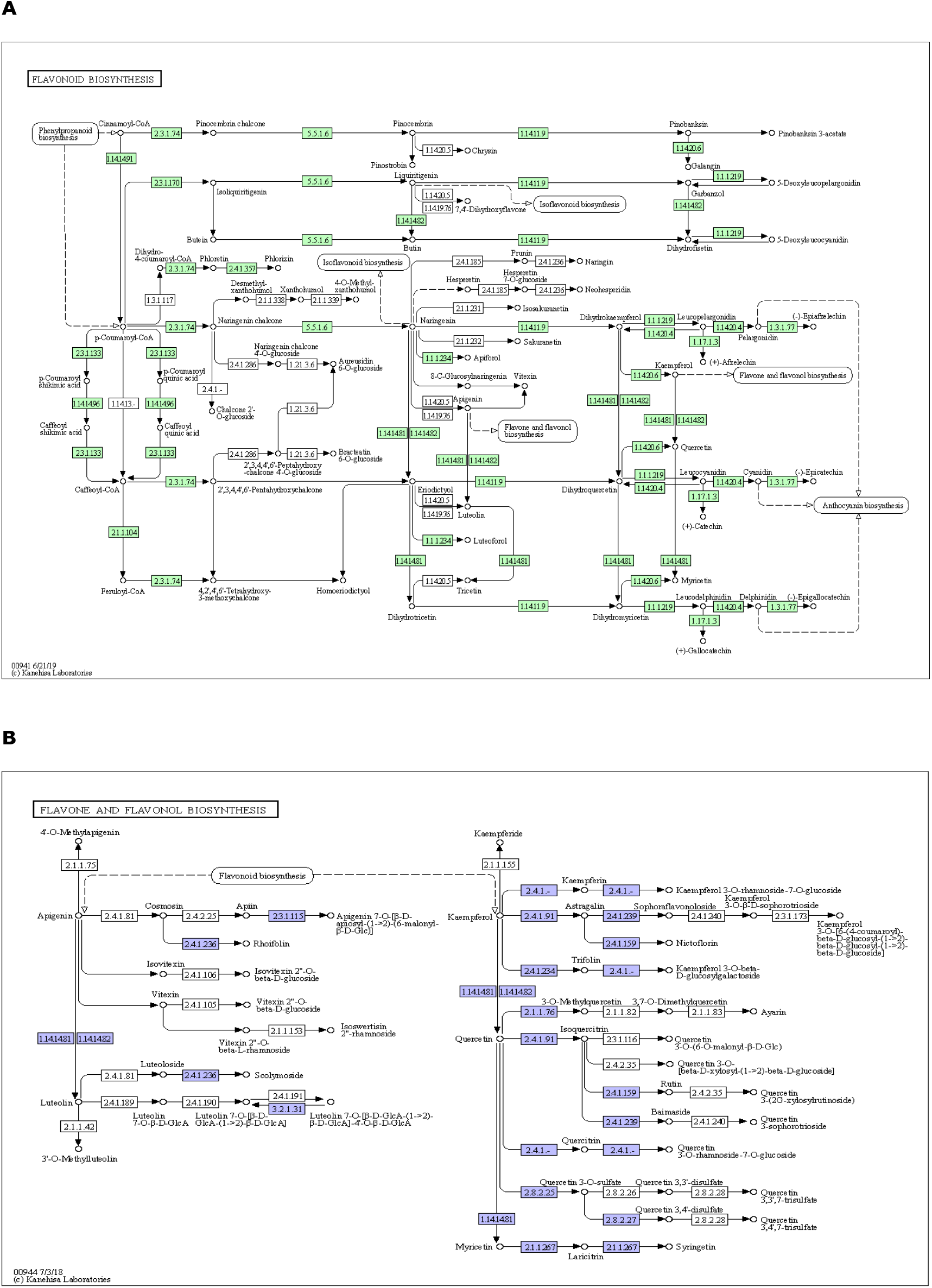
The flavonoid (A), and flavone and flavonol (B) KEGG bio-synthesis pathways in *Abelmoschus esculentus* (http://www.kegg.jp/kegg/kegg1.html). Putative okra enzyme coding genes for which a bi-directional best hit was found to enzymes pathways are shown with coloured EC identifiers.

## Supplementary tables

**Table S1.**
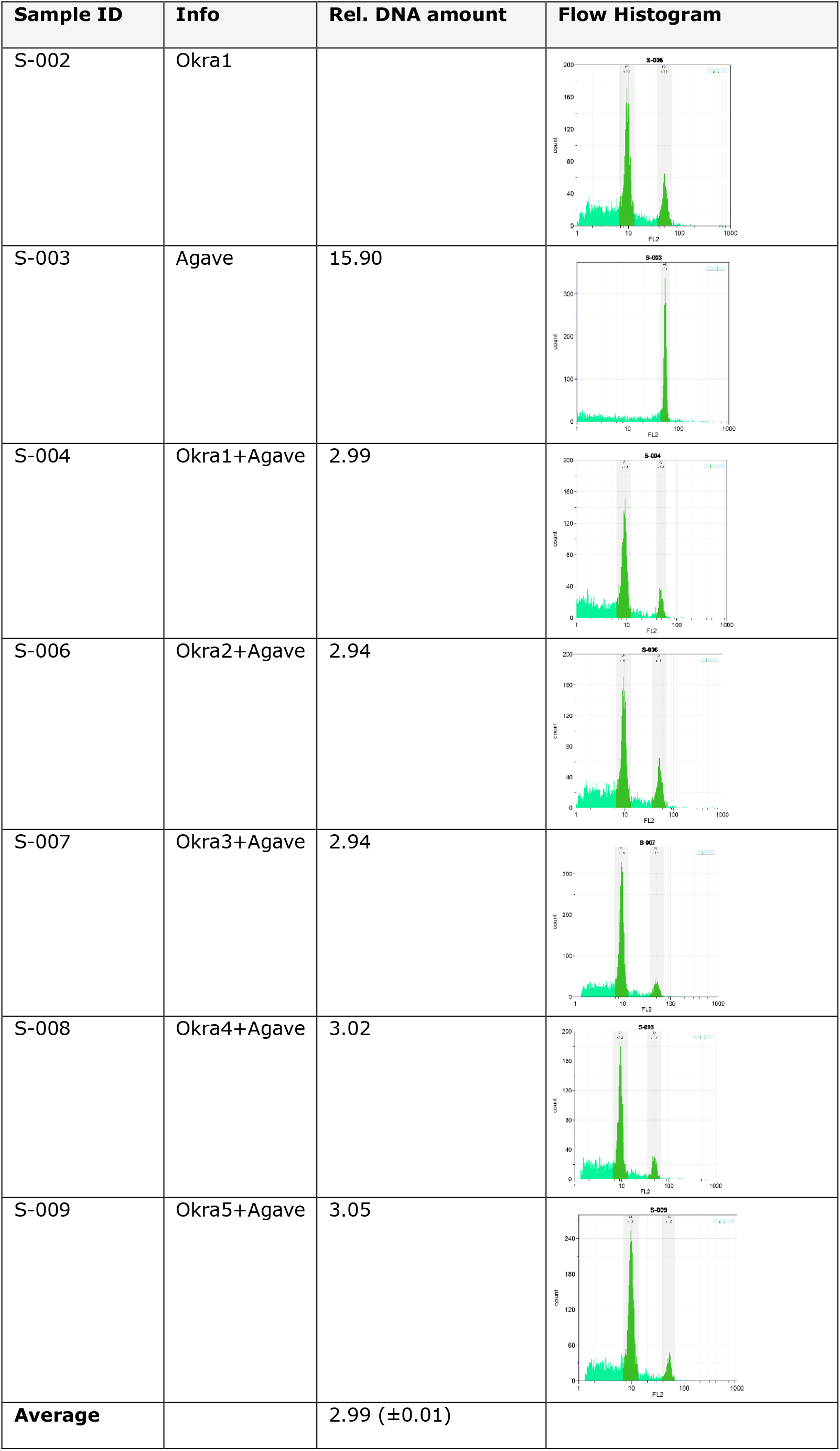
DNA amount of nuclei samples from okra root tip cells. DNA amount of okra replicate samples in picogram quantities was compared to a reference sample from *Agave Americana*. In the right column flow histograms of Okra samples and the Agave reference sample are shown. The count of observed nuclei in each histogram is depicted on the y-axis and is proportional to fluorescent intensity of each peak. The position of the peak along the x-axis is proportional to the relative DNA amount in each nuclei.

**Table S2.**
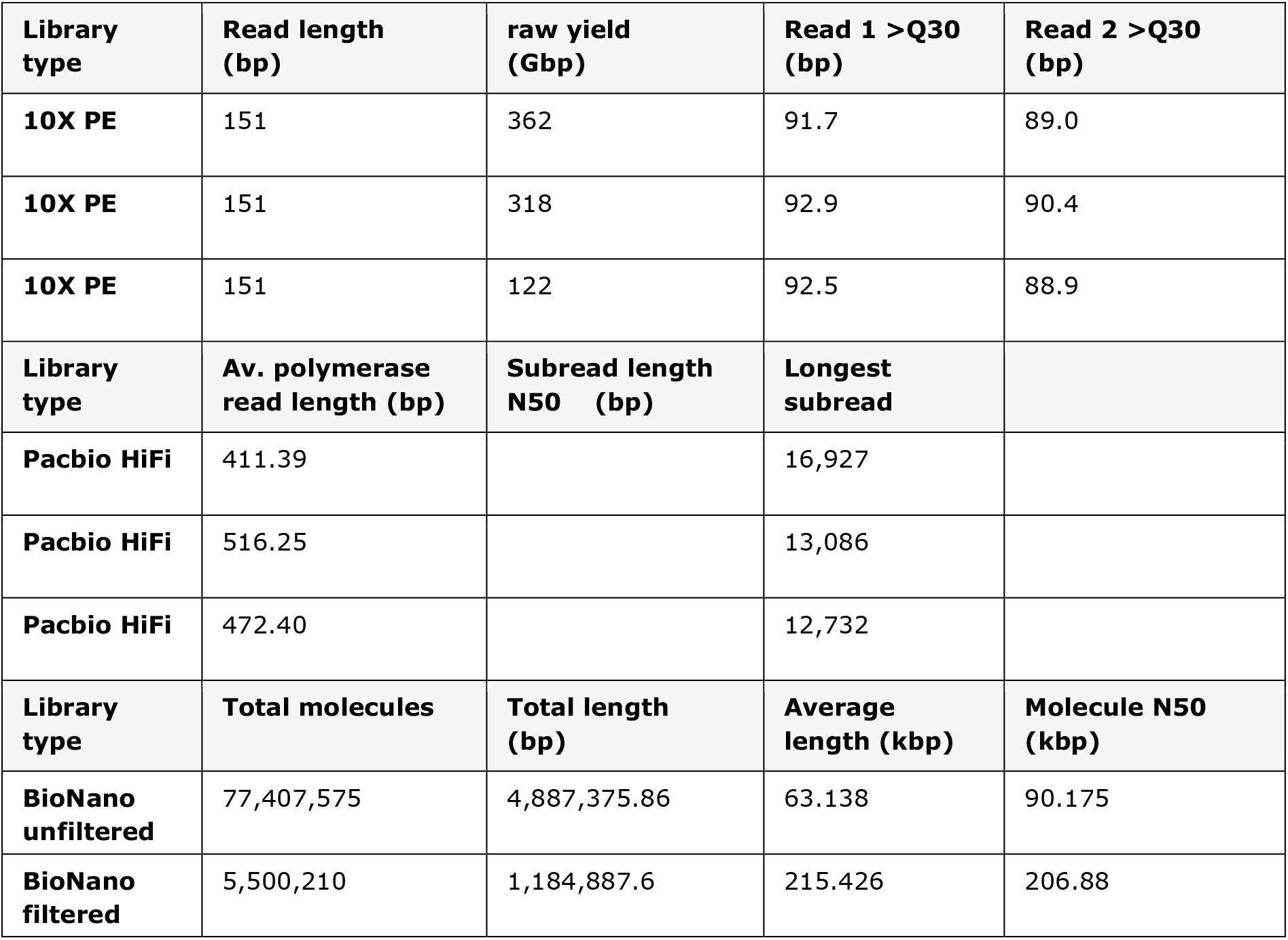
Genome sequencing and genome map data statistics. Linked read sequencing for three 10X Genomics libraries was performed using Illumina paired-end (PE) sequencing. Circular consensus sequencing (CCS) was performed for three Pacbio Hifi sequence libraries. Genome map data was produced for one BioNano DLE labelled library.

**Table S3.**
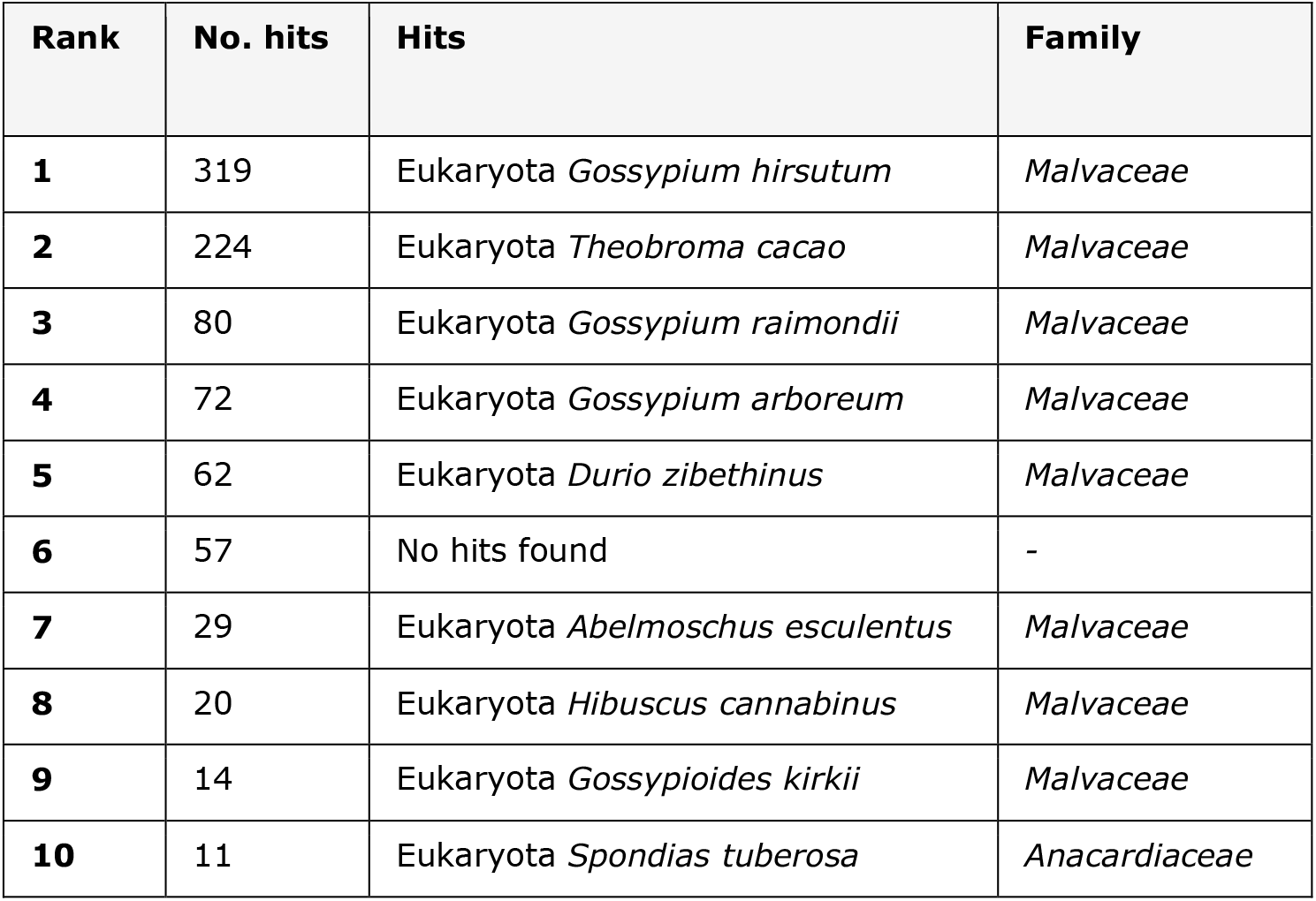
BlastN screening statistics for Pacbio HiFi reads. Screening was performed against the NCBI nucleotide database, Readouts are indicated in counts of *Malvaceae* species specific and non-*Malvaceae* hits for a subset of 1000 HiFi reads.

**Table S4.**
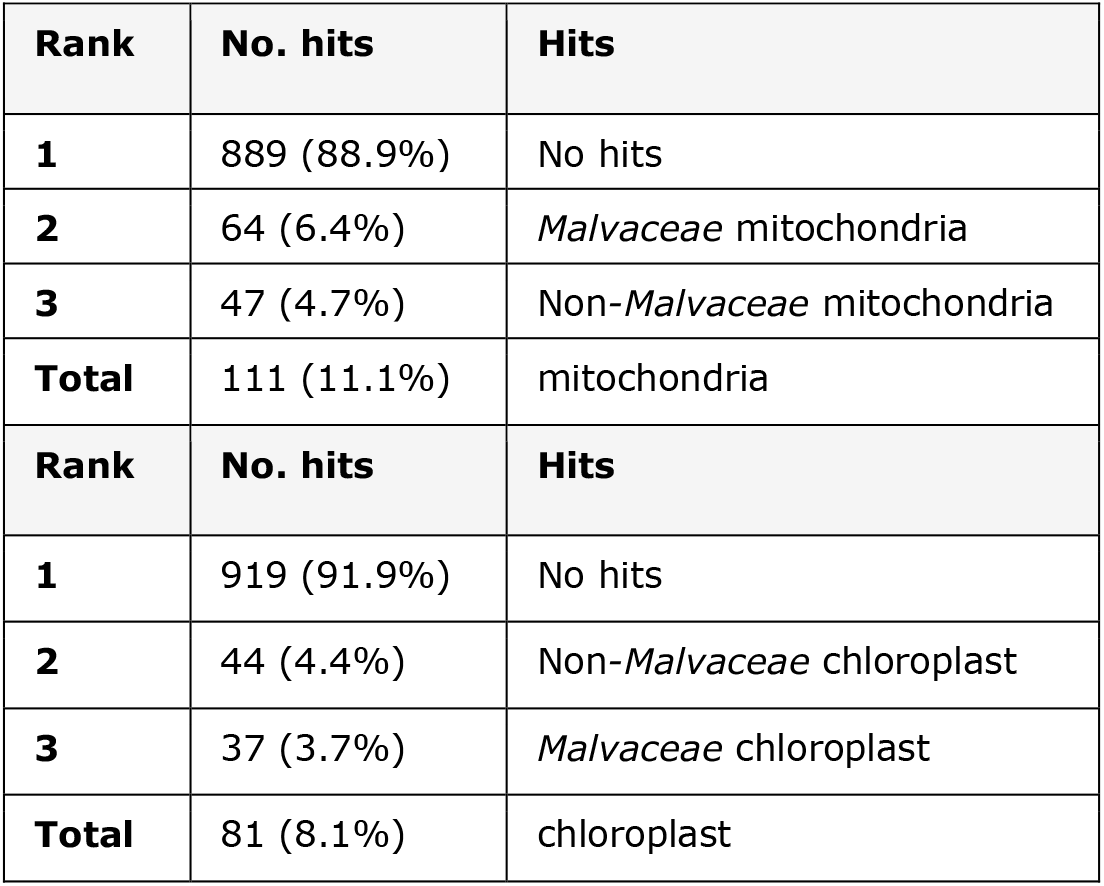
Pacbio sequence library contamination statistics for 1000 HiFi. Organelle content was determined using a BlastN screening against mitochondrial and chloroplast databases.

**Table S5.**
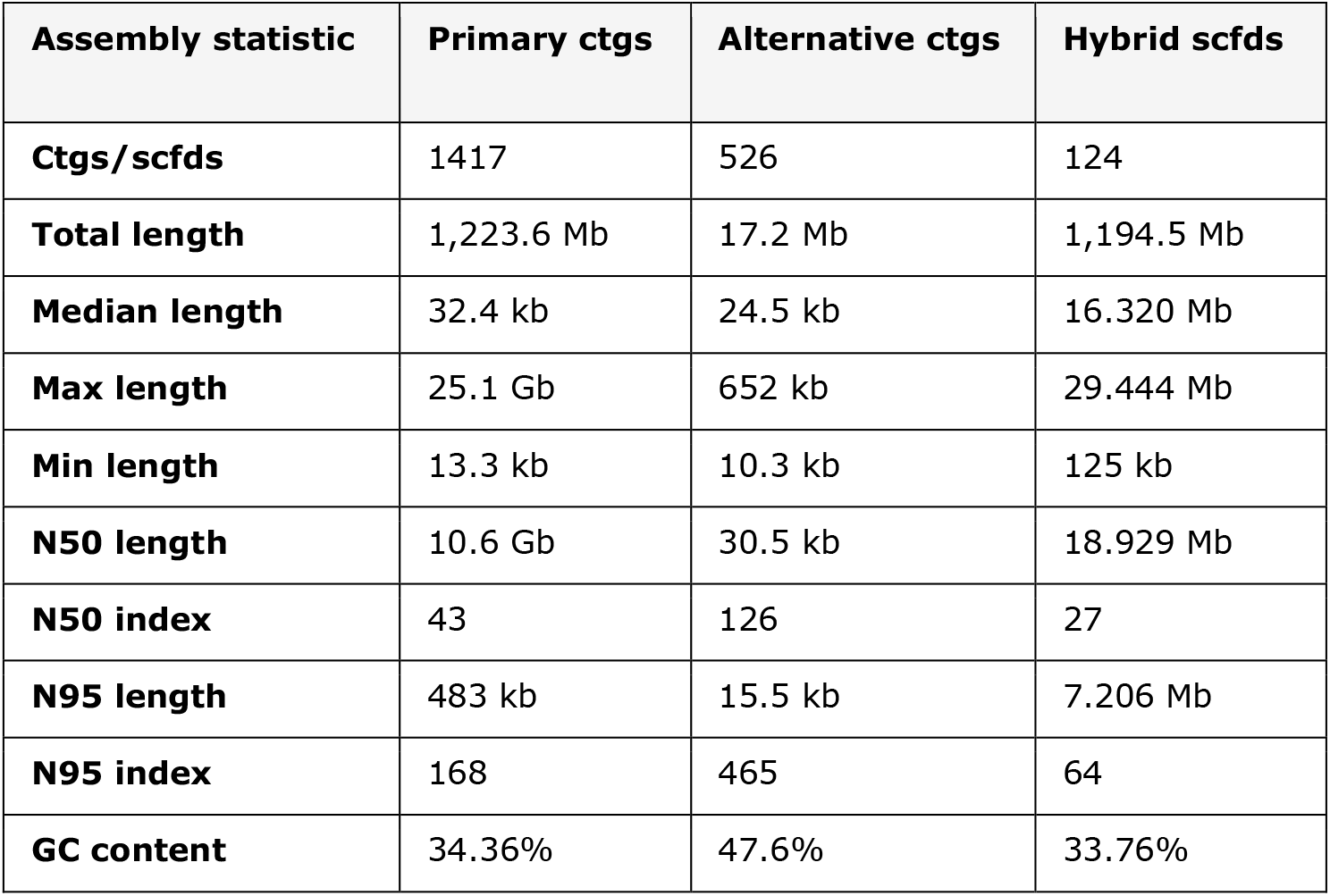
NGS assembly and hybrid scaffolding statistics. Sequences were assembled using the Hifiasm assembler and scaffold with Bionano Genomics genome maps.

**Table S6.**
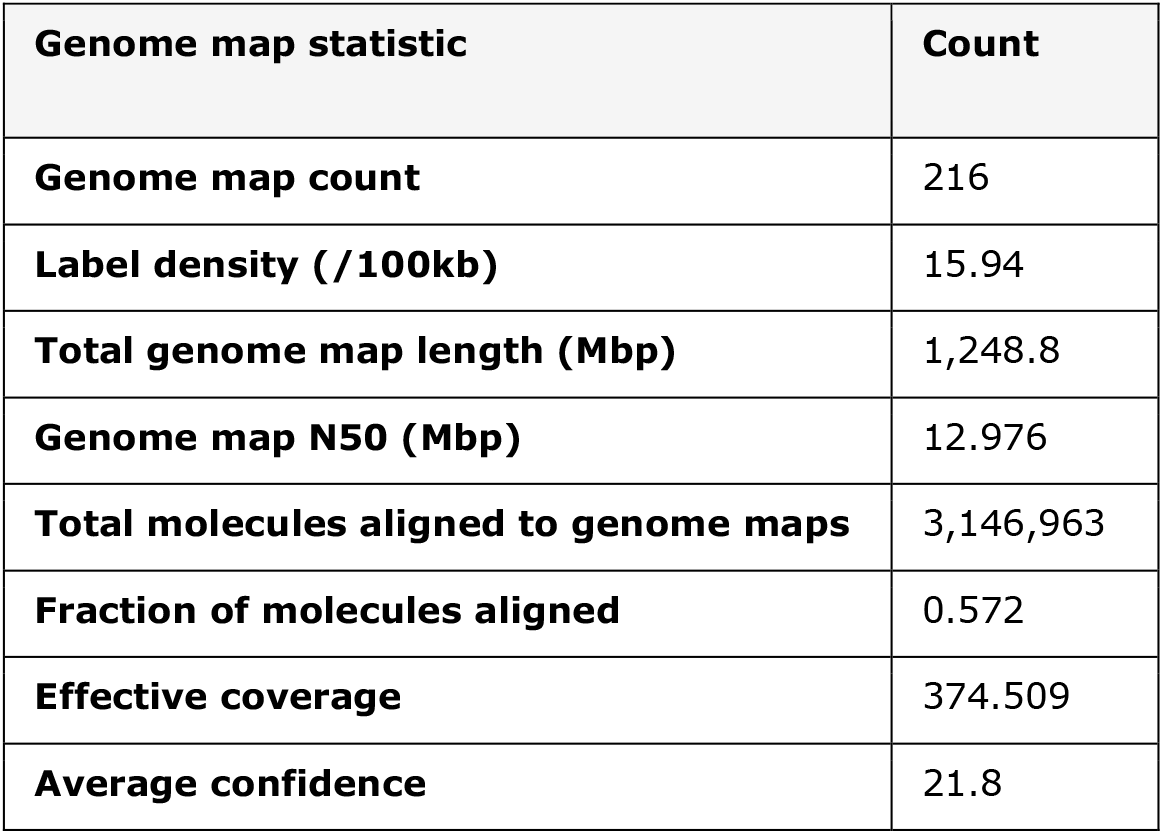
*De novo* genome map assembly statistics. Assembled molecules were mapped back to genome maps to estimate the effective coverage and average confidence the *de novo* assembly.

**Table S7.**
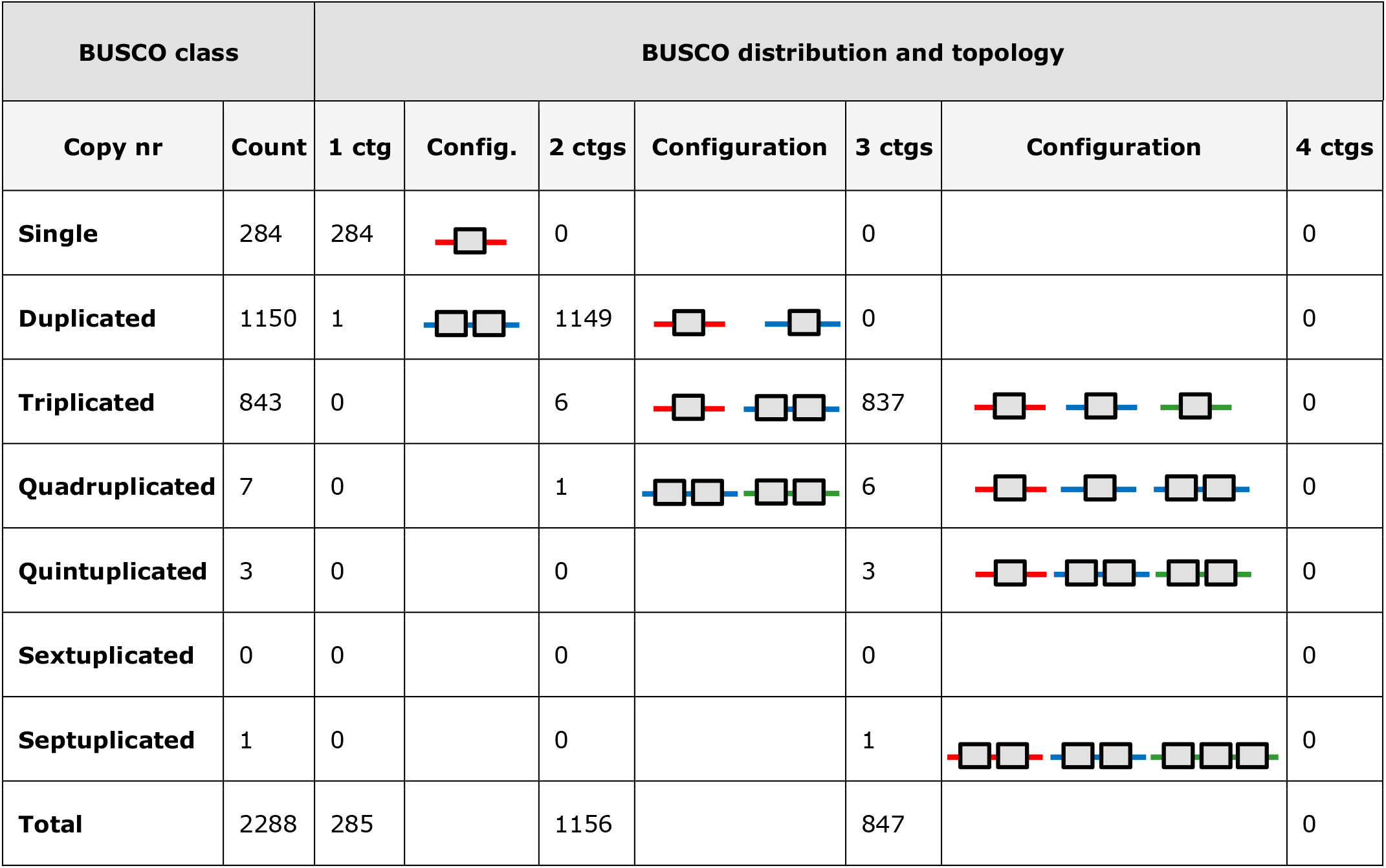
BUSCO distribution and topology. BUSCO genes are classified according to their copy number in the genome. Distribution counts for ortholog gene copies have been indicated according to their position either on one, two or three contigs. Configuration of gene copies is depicted by a horizontal line representing a contig and superimposed small grey coloured boxes representing a gene copy.

**Table S8.**
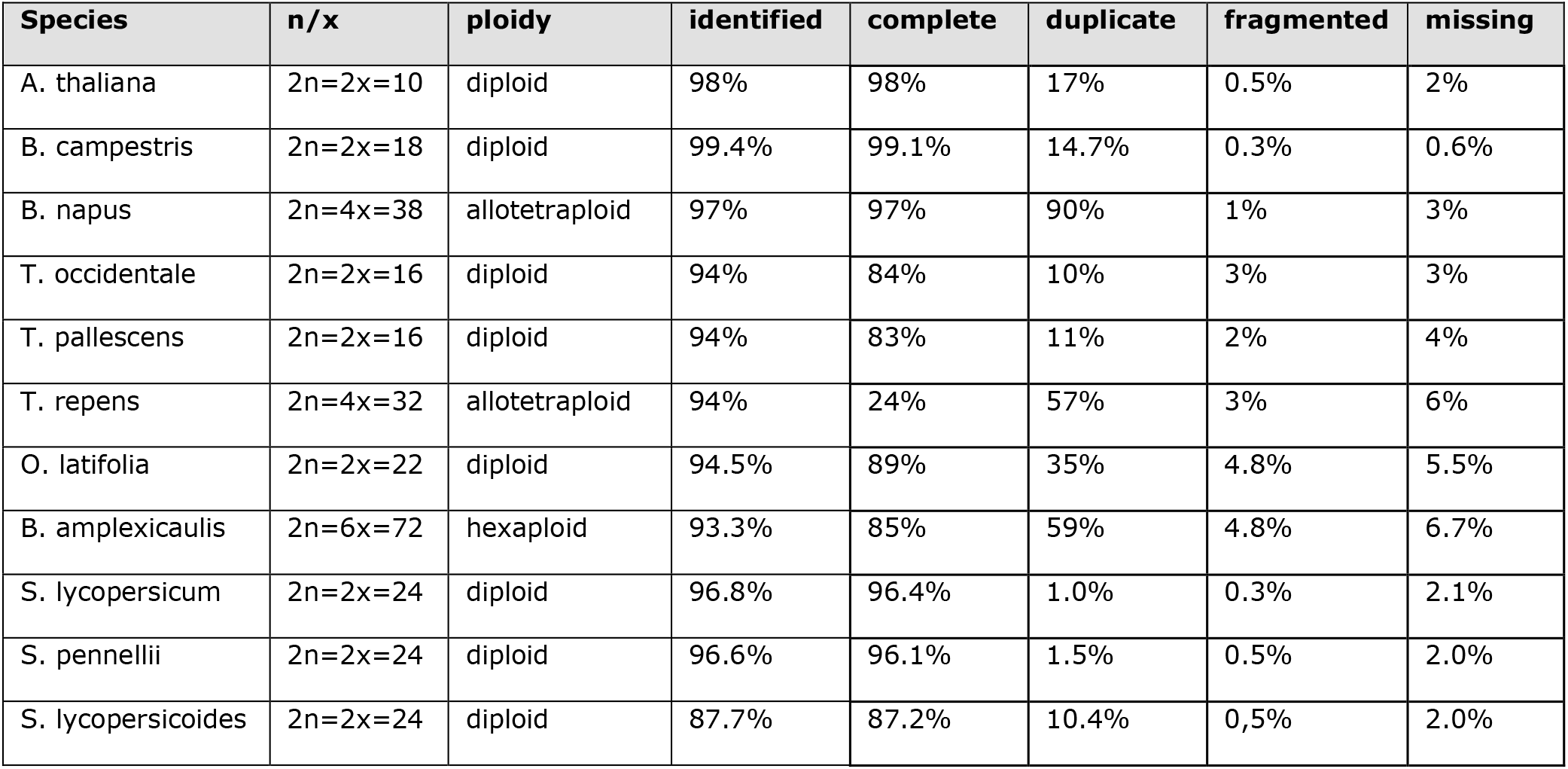
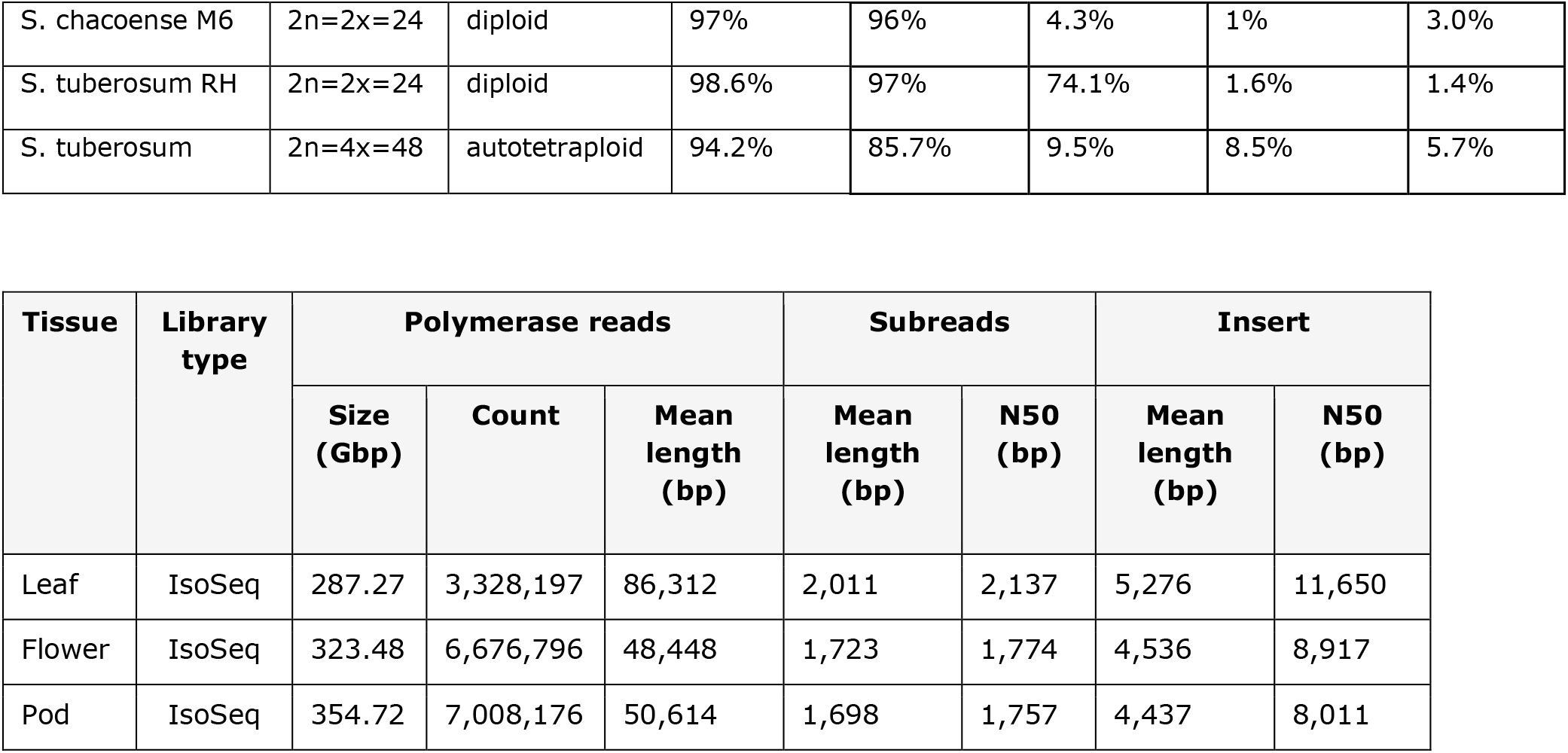
Transcriptome sequencing statistics. Pacbio IsoSeq libraries were constructed for 3 different tissues as indicated.

**Table S9.**
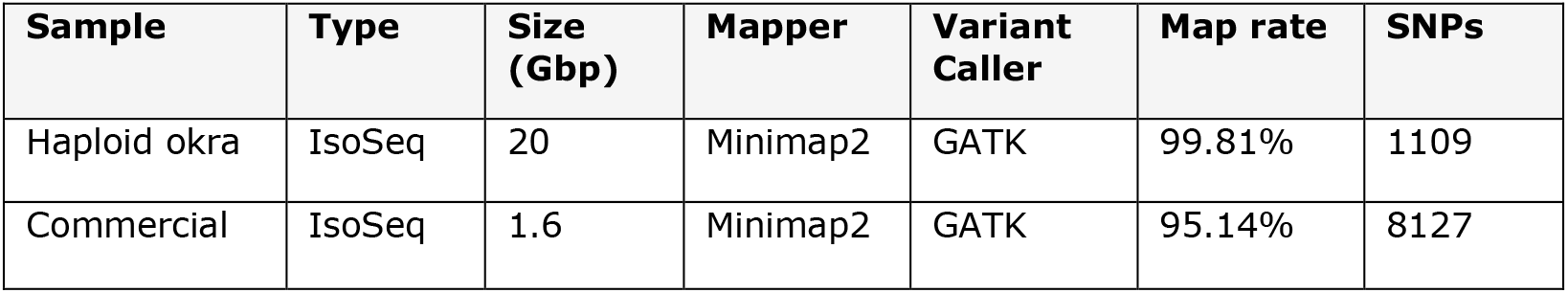
Transcriptome mapping to the okra reference genome and SNP calls.

**Table S10.**
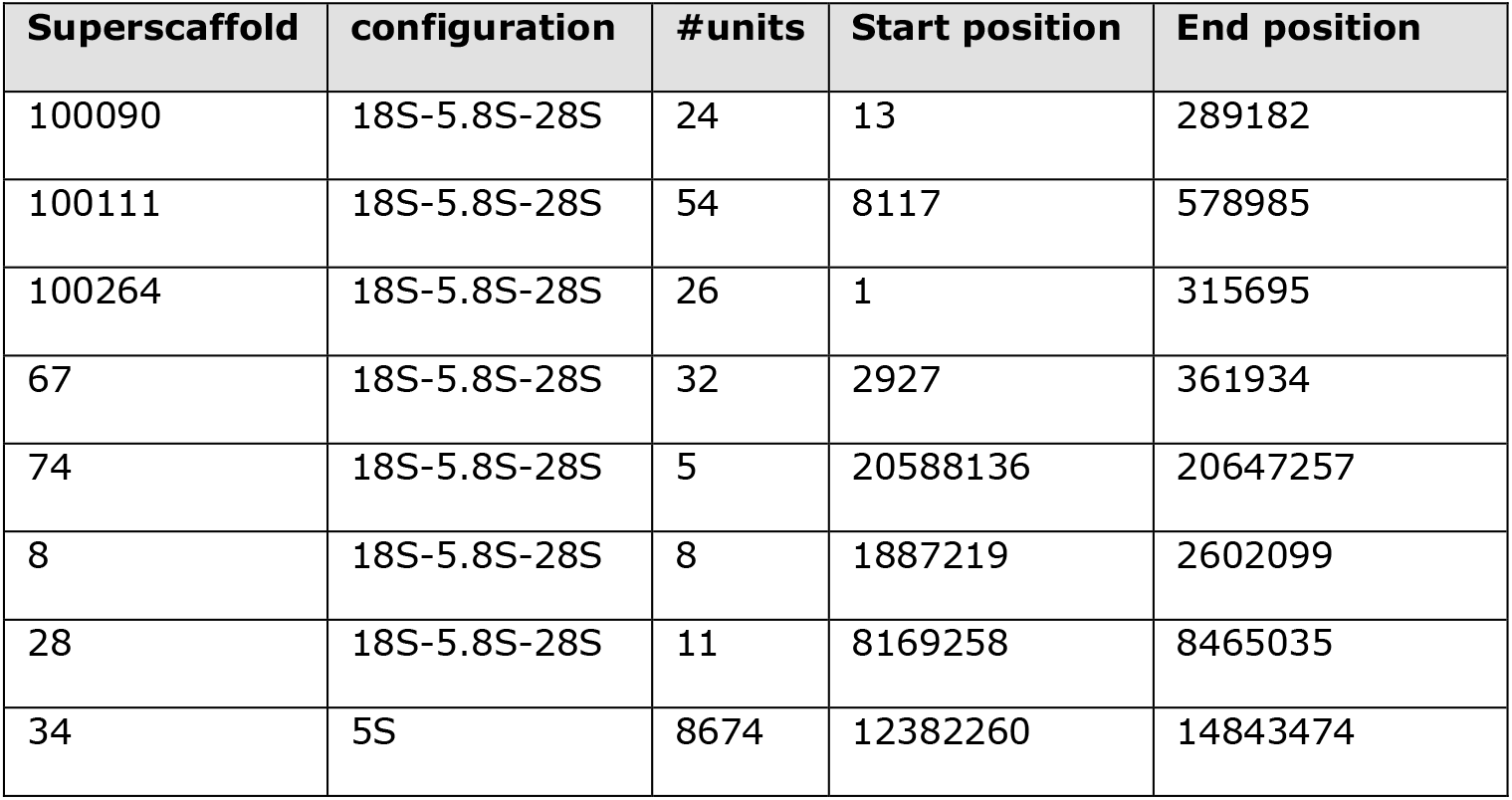
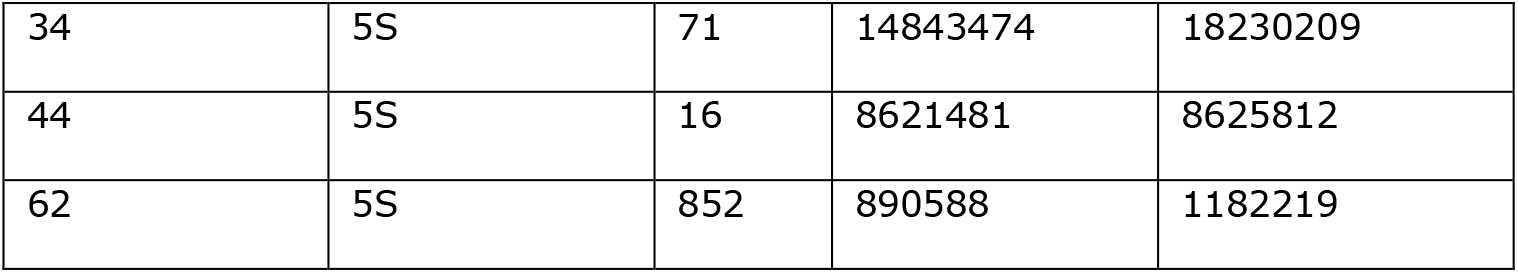
Ribosomal gene clusters in the okra genome. Ribosomal gene clusters are characterized by unit configuration and number of tandemly arranged unit copies per superscaffold. Total lengths of clustered units that can be derived from the start and end position.

## Materials and Methods

### Chromosome analysis

Plants of the Green Star F1 hybrid of okra (*Abelmoschus esculentus*) were grown in small for collecting actively growing rootlets that appeared at the outside of the pot soil. The root tips were pretreated with 8-hydroxyquinolin and then fixed in freshly prepared glacial acetic acid: ethanol 96% (1:3) and one day later transferred to ethanol 70% for longer storage at 4 ºC. Young flower buds were collected from nurse fields in Kamphaeng Saen, Thailand, and directly fixed in acetic acid ethanol without pretreatment. Microscopic preparations of root tip mitoses and pollen mother cells at meiotic stages were prepared following pectolytic enzyme digestion of cell walls and acetic acid maceration and cell spreading following the protocol of Kantama *et al*. (2017). Air-dried slides were stained in 300 nM 4′,6-diamidino-2-phenylindole (DAPI) in Vectashield (Vector Laboratories) and studied under a Zeiss fluorescence microscope equipped with 1.4 N.A. objectives and appropriate epifluorescence filters for DAPI. The captured images were optimized for best contrast and brightness in Adobe Photoshop, and slightly sharpened with the Focus Magic (www.focusmagic.com) 2D deconvolution sharpening to remove excessive blurring of the DAPI fluorescence (Kantama *et al*., 2017).

### Bionano optical maps

Sequence-specific labelling of approximately 700 ng genomic DNA from okra cv. Green Star and subsequent backbone staining and DNA quantification for BioNano mapping was done using a Direct Label Enzyme (DLE-1, CTTAAG) according to the manufacturer protocol 30206F BioNano Prep Direct Label and Stain Protocol (https://bionanogenomics.com/wp-content/uploads/2018/04/30206-Bionano-Prep-Direct-Label-and-Stain-DLS-Protocol.pdf). Chip loading and real-time analysis was carried out on a BioNano Genomics Saphyr® analyser, using the green color channel on 3 flow cells, according to the manufacturer system guide protocol 30143C (https://bionanogenomics.com/wp-content/uploads/2017/10/30143-Saphyr-System-User-Guide.pdf). Using the DLE-1 enzyme, 1.18 Tbp of filtered DNA molecules with an average length of 215kb was produced, with a label density of 15.9/100kb and a molecule N50 of 207 kb. Subsequently, a *de novo* assembly was constructed using Bionano Access™ (v.3.2.1) and the non-haplotype aware assembly program without extend and split but with cutting of the complex multi-path regions (CMPR). Per the default settings, molecules < 150 Kbp were removed before assembly. Next, a hybrid scaffolding of assembled sequence contigs was performed with Bionano Genomics Solve (v.3.2.1) with a 375X-fold coverage for the DLE-1 molecules. Molecule quality hybrid scaffold report were carried out using the BioNano Solve™ analysis pipeline (https://bionanogenomics.com/support-page/data-analysis-documentation/).

### Pacbio HiFi, linked-read sequencing and de novo assembly

We produced 3 Pacbio HiFi libraries using gDNA isolated from okra leaf tissue according to the manufacturers protocol (https://www.pacb.com). HiFi reads of 15-20 kb were generated by Circular Consensus Sequencing, using 6 SMRT cells, in total yielding 1,400 Gbp of sequence data. Subsequent consensus calling was done using the pbccs v5.0.0 command line utility. HiFi reads were defined as CCS reads having a minimum number of 3 passes and a mean read quality score of Q20. Reads from different libraries were then combined into a single dataset for further analyses. Assembly of HiFi Reads was done using hifiasm v0.12-r304 for coverages of ∼20X, ∼84X and ∼95X (Cheng *et al*., 2020). Primary contigs of the ∼95X coverage assembly were scaffolded using Bionano genomics Solve v3.6_09252020 and an optical *de novo* assembly. Solve scaffolded output was further scaffolded, in contrast to the unscaffolded output, using Arcs v1.2.2 (https://github.com/bcgsc/arcs) and Links v1.8.7 (https://github.com/bcgsc/LINKS) based on the 10X genomics data that was mapped using Longranger v2.2.2. Scaffolds resulting from the final step were renamed to fit the naming scheme from the Bionano scaffolding.

The 10X Genomics libraries were constructed with the Chromium™ Genome Reagent Kits v2 (10X Genomics®) according to the Chromium™ Genome v2 Protocol (CG00043) as described by the manufacturer (https://www.10xgenomics.com). 10X Genomics libraries were sequenced on 2 separate runs using the Illumina Novaseq6000 platform and S2 flow cells. Base calling and initial quality filtering of raw sequencing data was done using bcl2fastq v2.20.0.422 using default settings. The Long Ranger pipeline from 10X Genomics was used to process the 800 Gbp sequencing output and align the reads to the tomato reference genome. After detecting the conflict region with the Bionano Genomics Access Suite (v.1.3.0), we manually inspected the conflict regions using 10X linked-reads mapped to the superscaffolds. Mapping and visualization of scaffolds was done with Longranger WGA v.2.2.2 and Loupe v.2.1.1 respectively.

### Assembly QC

Statistics on N*X* lengths, GC-percentage, mean-, median-, maximum- and minimum lengths of contigs or scaffolds for each assembly step was collected from software output and when not available was generated with custom python scripts. HiFi reads were mapped back to the assembly using minimap2 v2.17-r941 to assess purging correctness with purge_haplotigs (Li, 2018; Roach *et al*., 2018). Minimap2 alignment was also used for blobtools v1.1.1 analysis to check the taxonomic origin, coverage and GC-percentage of unscaffolded output (Laetsch *et al*., 2017). The completeness of the assembly was benchmarked using BUSCO with the eudicots_odb10 (eukaryota, 2020-09-10) lineage set to scan for single copy orthologs (Kriventseva *et al*., 2019; Simão *et al*., 2015). Augustus v3.2.2 was subsequently used for gene prediction. BUSCO output was used for topology analysis of duplicated genes (Stanke *et al*., 2006). To assess repeat content and synteny within the scaffolds, the assembly was self-aligned using a combination of minimap2 and Dgenies, and nucmer v4.0.0beta2 together with mummerplot v3.5 (Kurtz *et al*., 2004).

### Iso-Seq sequencing and data analysis

Total RNA was isolated from leaf (10 µg), flower buds (21 µg) and young fruits (31µg) from okra cv. “Green Star”. RNA quality was checked on a Bioanalyzer platform (https://www.agilent.com) by comparing to standard samples of 25S and 18S ribosomal RNA. Transcript samples were subsequently used for construction of 3 sequence libraries and sequenced with PacBio SMRT technology (https://www.pacb.com/smrt-science/smrt-sequencing) using 4 SMRT cells. Consensus reads were produced with the ccs v5.0.0 command-line utility of PacBio. HiFi reads were classified as such using the same specifications as the genomic reads. Primers sequences from reads were removed and demultiplexed with lima v2.0.0. Poly-A tails were trimmed and concatemer were removed with isoseq3 v3.4.0 refine to generate full length non-concatemer reads and subsequently clustered with isoseq3 cluster. Since distributions of mean read quality showed over 90% of data to have a mean quality score in range of 90-93, no final polishing was applied. High quality full-length transcripts obtained from the SMRT analysis pipeline were then mapped to the hybrid assembly using Gmap (Wu and Watanabe, 2005).

### Read analysis

Quality control of reads was performed using SMRTlink v9 and FastQC v0.11.9. A random sample of 1000 reads per SMRT cell was taken using seqtk v1.3-r106 seq with a random seed from the bash v4.2.46 internal pseudorandom generator. Samples were screened for chloroplast content, plastid content and taxonomic contamination applying BLAST v2.10.1+ with parameter settings *-evalue 0*.*001* and *-max_target_seqs 1* (Altschul *et al*., 1990; Altschul *et al*., 1997; Camacho *et al*., 2009). The databases used for each screening were NCBI nt, plastid and mitochondrion publicly available FTP downloads dated 2020-11-15 (Agarwala *et al*., 2016).

K-mers were counted for both 10X linked reads and HiFi reads, using kmc 3.1.1 (Kokot *et al*., 2017) with parameter settings -*m64-ci1-cs10000* for *k = 16, 21, 28, 37, 48* and *61* to determine the k-mer size for best-model fit. For 10X linked reads 23 bp of R1, containing 16 bp barcode plus the 7 bp long spacer sequence, were trimmed off before counting. The kmc histogram of counts was subsequently used for estimation of genome parameters and visualization of k-mer spectrum using genomescope2 (Ranallo-Benavidez *et al*., 2019). The polyploid nature of okra was further examined applying a locally developed, publicly available fork of smudgeplot labeled v0.2.3dev_rn that is true to the original algorithm, allowing for parallelization. In the original algorithm k-mer pairs with a hamming distance of 1 are found by a recursive method that iterates over all positions within the k-mer. The redesigned algorithm parallelizes the search by each thread looking at a given position within the k-mer. To reduce the number of false negatives, results are then filtered using a bloom filter with an *error rate* set at *0*.*0001*.

### Annotation

Repeats annotation was carried out with RepeatModeler, RepeatClassifier and RepeatMasker tools and the combined RepBase (2014) and Dfam (2020) databases for classification of repeats (Bao *et al*. 2015; Hubley *et al*., 2016; Smith *et al*., 2013). Full length non-concatemer IsoSeq reads were mapped against the genome assembly using minimap2 with parameter settings *-ax splice -uf --secondary=no -C5*. A repeat masked genome and transcriptome read mapping was then input to the Braker2 v2.1.5 pipeline to generate a structural annotation of genes using *ab initio* prediction (Borodovsky and Lomsadze, 2011; Hoff *et al*., 2019). Using the general feature format (GFF) file output, open reading frame translations of the predicted genes were made with the default eukaryotic translation table. Produced polypeptide sequences were then annotated with interproscan v5.39-77 10 that collected annotations from the default set of databases including Panther, Gen3D, CDD, etc (Hunter *et al*., 2009; Jones *et al*., 2014).

### Variant calling

To investigate the diversity among okra accessions, multiple publicly available transcriptome datasets were mapped to the assembled okra reference as presented in this study. Public datasets include SRR620228 (https://www.ncbi.nlm.nih.gov/sra/?term=SRR620228), PRJNA393599 (https://www.ncbi.nlm.nih.gov/sra/?term=PRJNA393599 and PRJNA430490 (https://www.ncbi.nlm.nih.gov/sra/?term=PRJNA430490). In case multiple tissues or runs were available, data were merged. Mapping was done using STAR v2.7.8a (Dobin *et al*., 2013; Leinonen *et al*., 2011). Duplicates were marked applying GATK v4.2.0 MarkDuplicatesSpark mode (McKenna *et al*., 2010). Then variants were called using GATK HaplotypeCaller with default filters. Obtained variants were then filtered for SNPs and subsequently quality filtered by GATK SelectVARIANTS with parameter filter settings *QD > 2*.*0 && MQ > 40*.*0 && FS < 60*.*0 && SOR < 3*.*0 && QUAL > 30*.*0 && MQRankSum > -12*.*5 && ReadPosRankSum > -8*.*0*. IsoSeq reads were mapped as described above.

### Subgenome separation

To separate the subgenomes in the okra genome assembly, a clustering approach on the repetitive part of the okra genome was applied to generate clusters of scaffolds with similar repeat patterns. For each scaffold, forward and reverse complement non-canonical k-mers were counted setting a length *k* = 13. The counts were normalized by scaffold length and then filtered to have a minimal count of 100. Sets of k-mer counts for both forward and reverse strand of each scaffold were then merged. All counts were subsequently increased by one and log10 transformed to generate a scale that was appropriate for visualization. Euclidean distances between the scaffolds were stored and scaffolds were subsequently clustered using the Ward’s (minimum variance) method. To visualize the clustering along the k-mer patterns, we generated a cluster map from a heatmap for the k-mer counts combined with a hierarchical clustering of the scaffolds. In addition, the clustering of kmers generated from an in-house generated artificial potato tomato hybrid assembly, and the allopolyploid cotton genome was used as a test set. Clusters were then compared based on BUSCO score completeness and scanned for homoeology in a network visualization using an adjacency matrix. Scores in the adjacency matrix were taken from the edge counts between scaffolds that have identical BUSCO genes in common, implying that scaffold x and scaffold y have an edge count z when sharing a number of z BUSCO genes. *Statistical analysis of metabolic pathway assignment for okra orthologs*

The mapping probability for okra orthologs to KEGG reference pathways was based on a hypergeometric test (one-sided Fisher’s exact test) to measure the statistical significance for pathway assignment of a putative okra ortholog set. Pathway *p*-values were calculated according to equation 1 (Eq. 1), where K equals the unique enzymes known for a pathway p, k for the number of searched enzymes uniquely mapping on pathway p, N as the number of unique enzymes of all reference species known for all pathways, and n the number of searched enzymes uniquely mapping on all pathways.

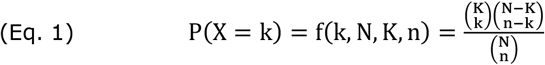

