## Supplementary material for "Genome and transcriptome architecture of allopolyploid okra *(Abelmoschus esculentus)*": Supplementary data_Profiling the Okra genome and transcriptome_BioRxiv.pdf

**Supplementary figures**

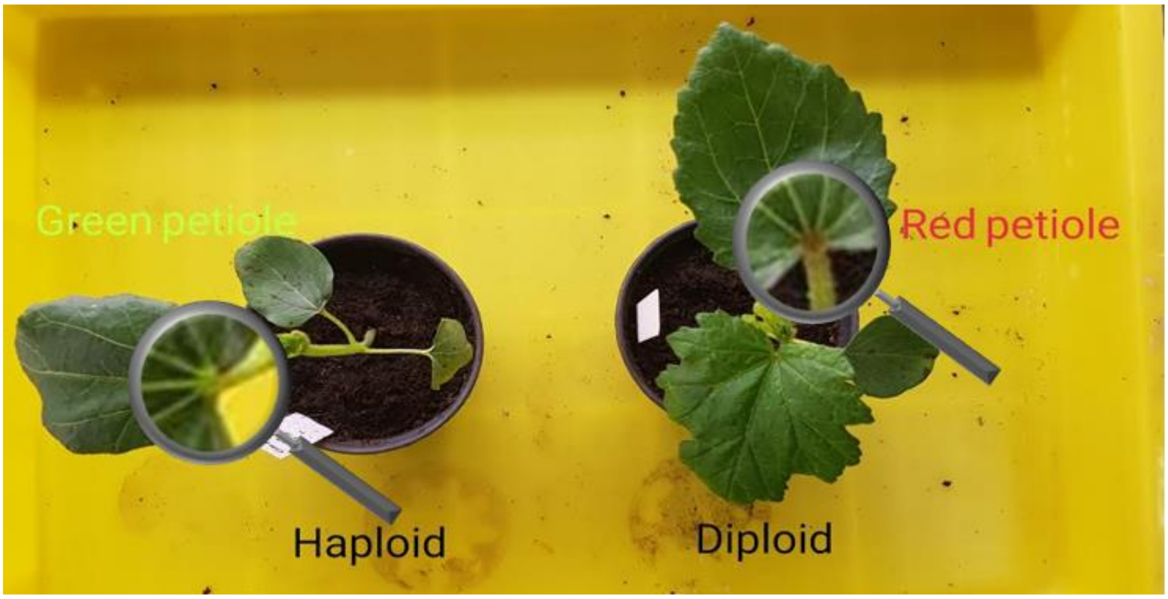


**Figure S1:** Phenotypes of diploid and haploid Okra plants. The magnifying glass in the image is placed over the position of the green and the red petiole for the haploid (left) and diploid (right) Okra plant respectively.

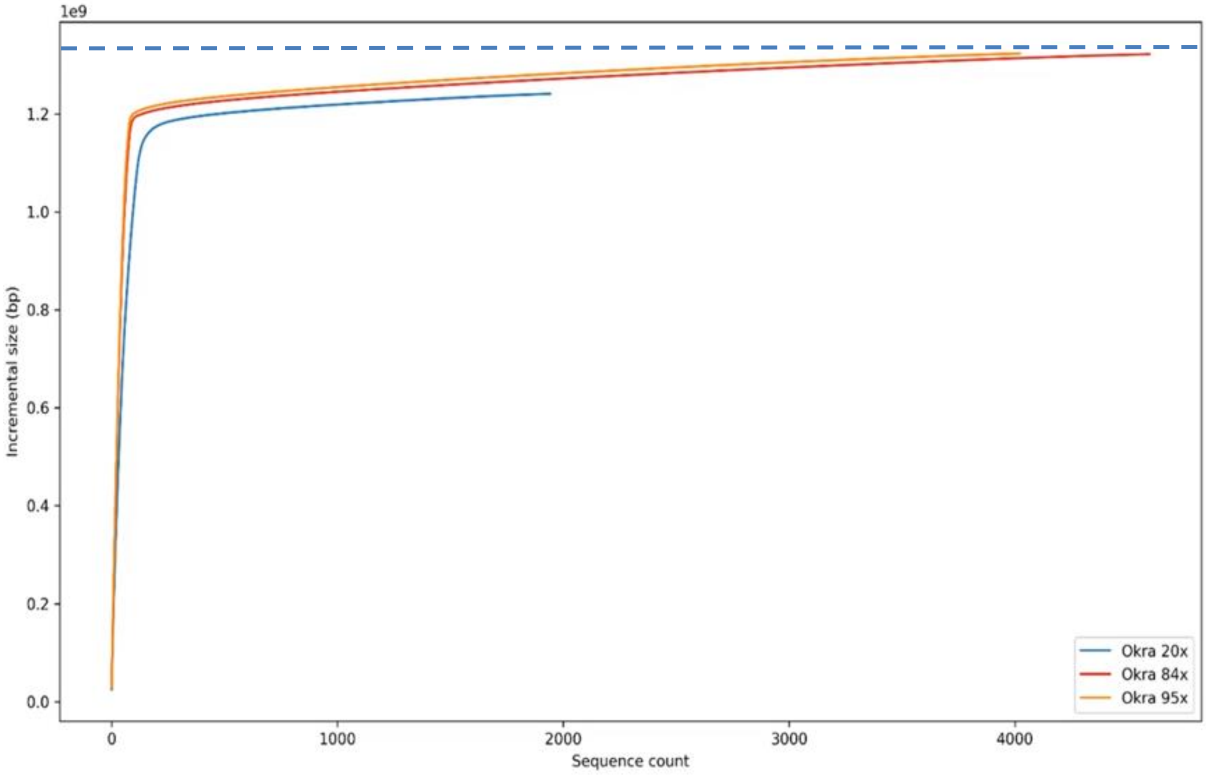


**Figure S2:** The incremental genome assembly size for Okra. The A50 plot for contigs larger than 100 bp shows the assembly size in Gbp on the y-axis is plotted against the incremental contig count at 20X, 84X and 95X sequence coverage indicated by the light blue, red and orange curve respectively.

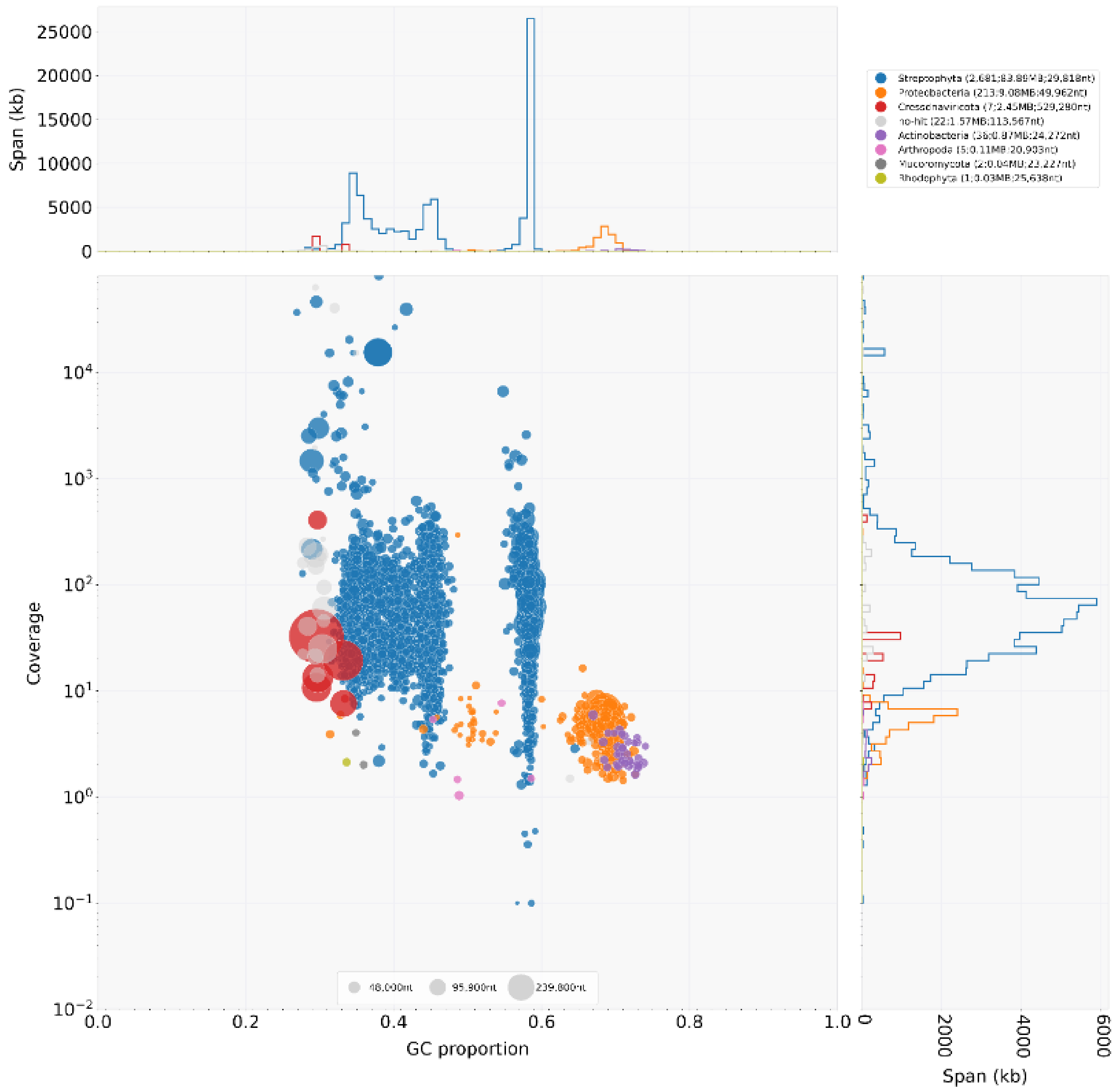


**Figure S3:** Taxon annotated GC coverage plot. In the low left panel the proportion of GC bases (x-axis) and read coverage (y-axis) for 527 alternative contigs are shown. Each coloured dot in the graph corresponds to a contig. Colours correspond to species classes for which a best BlastN match was found in annotated databases. In the top left graph and the low right graph the relative proportion for each class is depicted with respect to the coverage and GC content, for which colour codes match species classes as indicated in the legend at the top right.

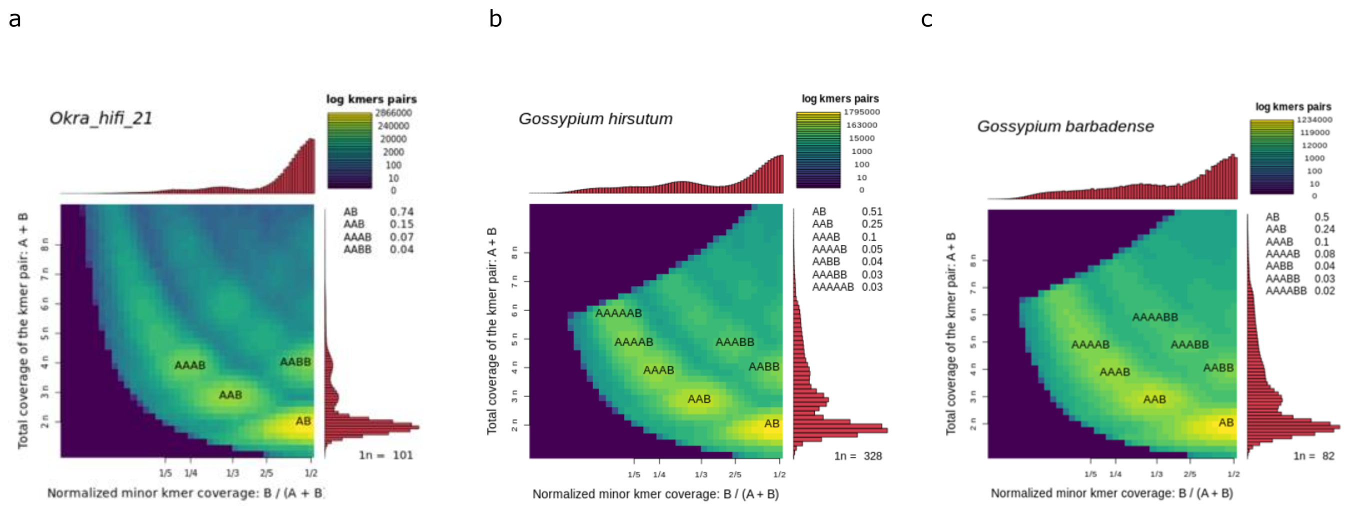


**Figure S4:** Smudgeplots for and haploid Okra (*Abelmoschus esculentus*) (a), and two allotetraploid cotton species *Gossypium hirsutum* (b), and *Gossypium barbense* (c). Smudgeplots are shown in log scale. The coloration indicating the approximate number of k-mer pairs per bin and the fraction of each kmer type is indicated in the top right legend of each plot.

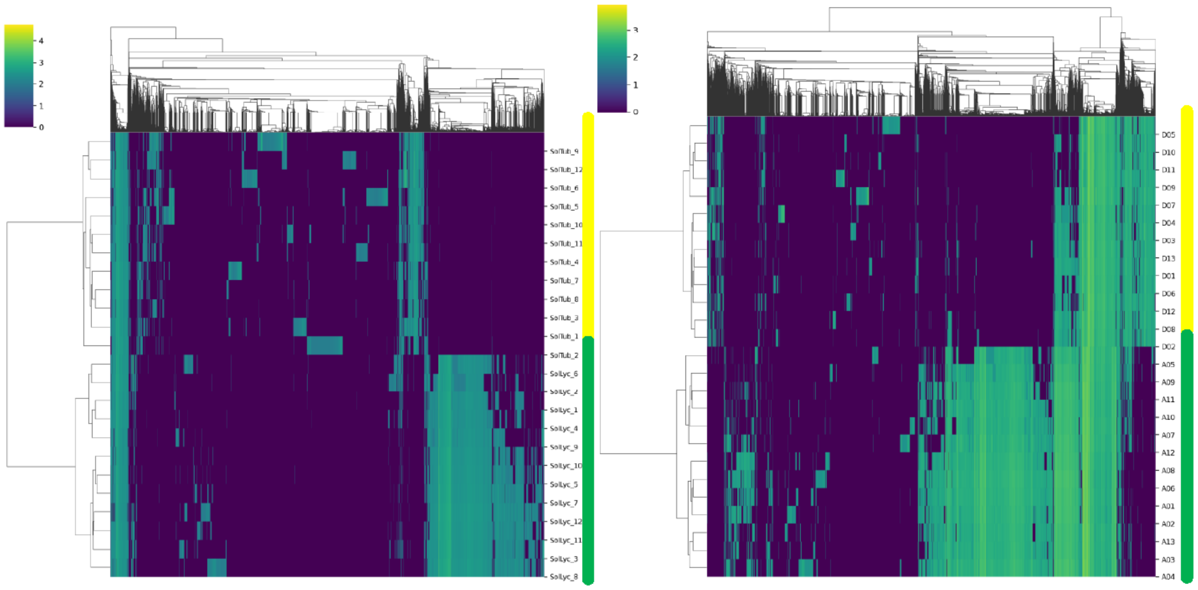

**Figure S5:** Cluster maps of repetitive 13-mer counts. K-mers generated for an artificial hybrid genome constructed from merged genomes of *S.lycopersicum* SL.4.0 and *S. tuberosum* cv. Solyntus (left panel) and from the allotetraploid cotton *G. hirsutum* genome. Yellow and green bars next to the chromosome identifiers mark the potato and tomato chromosomes (left panel) and the cotton chromosomes from the A and D subgenomes (right panel) respectively. Color code bar indicates log10 scaled kmer counts.

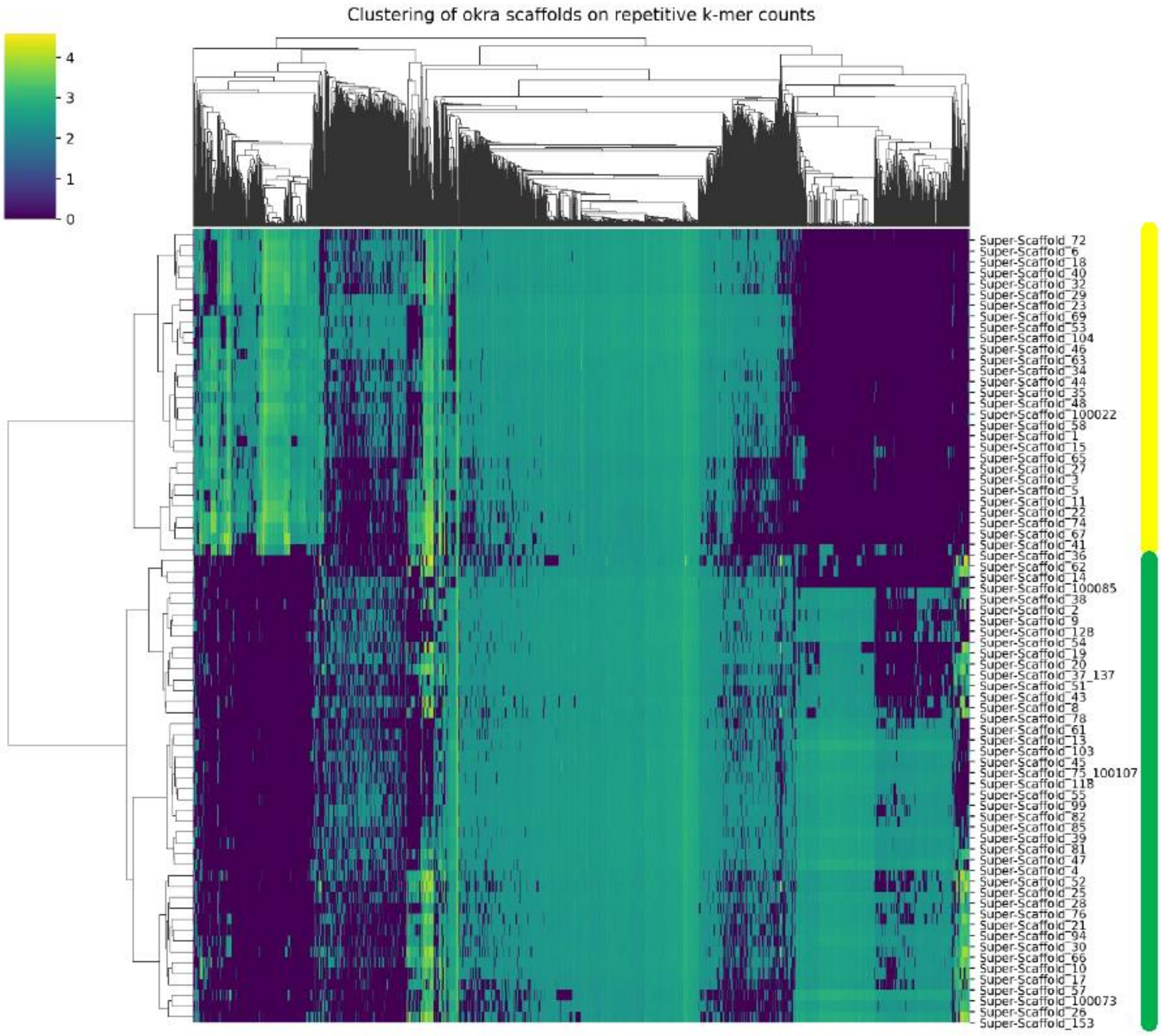


**Figure S6:** Cluster maps of repetitive 13-mer counts for the okra reference genome. Yellow and green bars next to the identifiers mark the superscaffolds clustering in cluster 1 and 2 respectively. Color code bar indicates log10 scaled kmer counts. A clear separation between cluster 1 and 2 based on repeat count and distinct repeat profile is apparent.

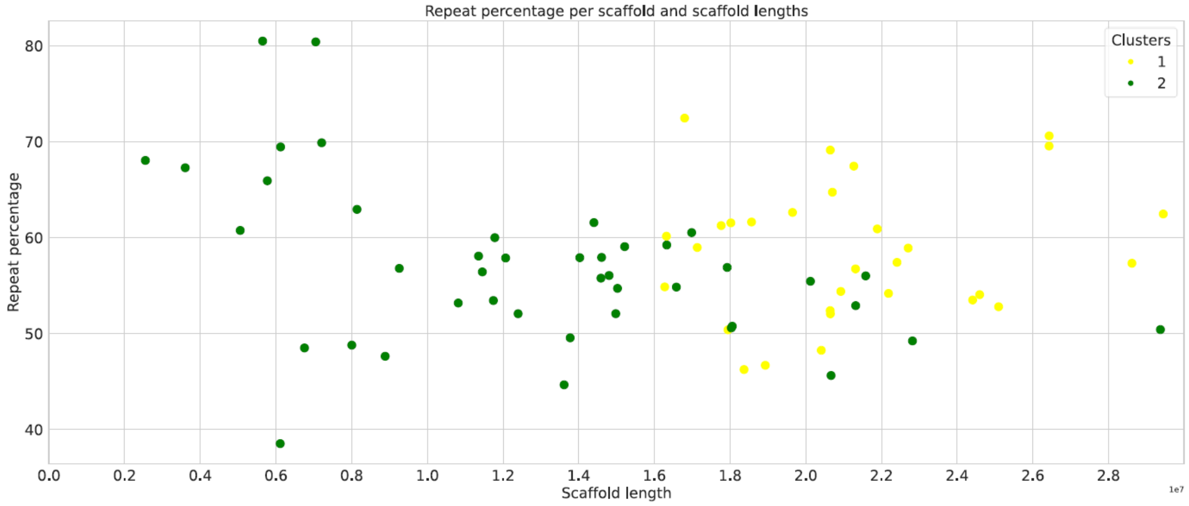


**Figure S7:** Repeat content analysis in superscaffolds. Superscaffolds assigned to cluster1 and 2 are represented by yellow and green dots respectively, and have been separated by scaffold length (x-axis) and repeat percentage (y-axis).

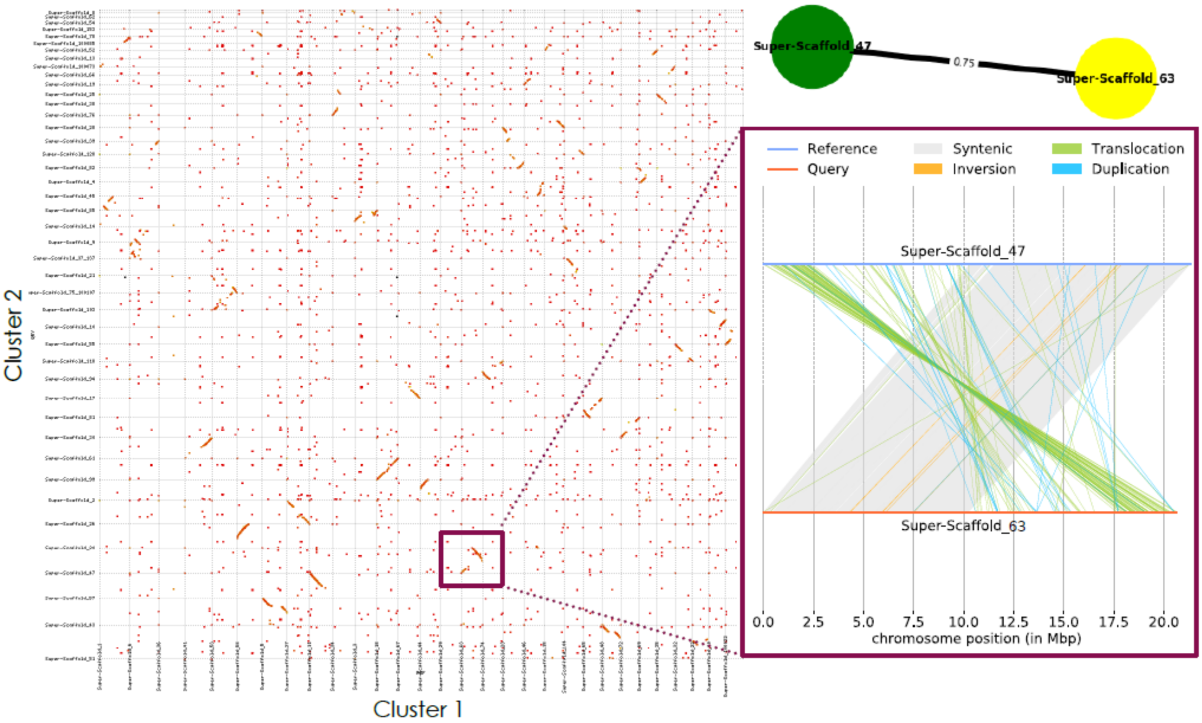


**Figure S8:** Dot plot alignment of superscaffolds from subgenomes. Superscaffolds are assigned to cluster 1 or 2 according to their kmer clustering profile. The top right graph shows two homoeologous superscaffolds 63 (cluster 1) and 47 (cluster 2) having 75% of BUSCO genes in common. The bottom right alignment detail of the aforementioned superscaffolds are partially syntenic, sharing a large inversion.

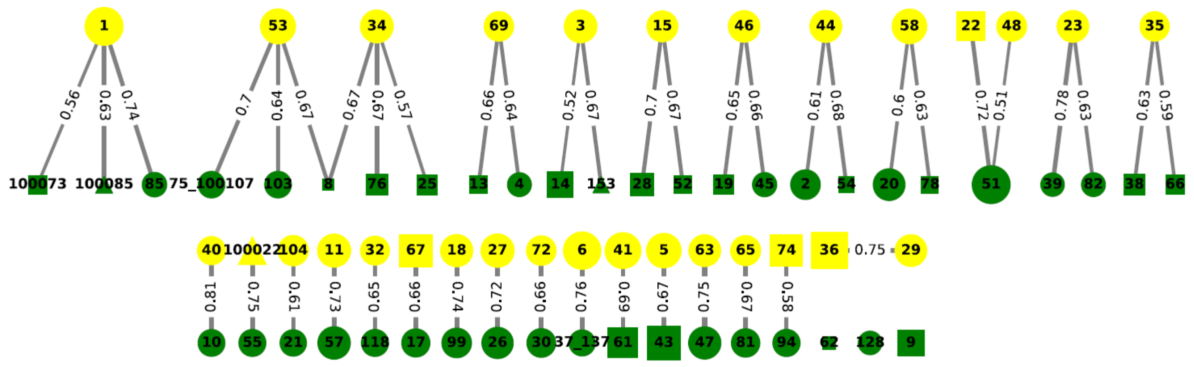

**Figure S9:** BUSCO connectivity graph. Yellow and green color coded nodes correspond to scaffolds from cluster 1 and 2 respectively. Edges between the nodes indicate the percentage of shared BUSCO genes between each scaffold pair. Note that a single node can have multiple edges. Pairs of scaffolds point to links of homoeology between scaffolds.

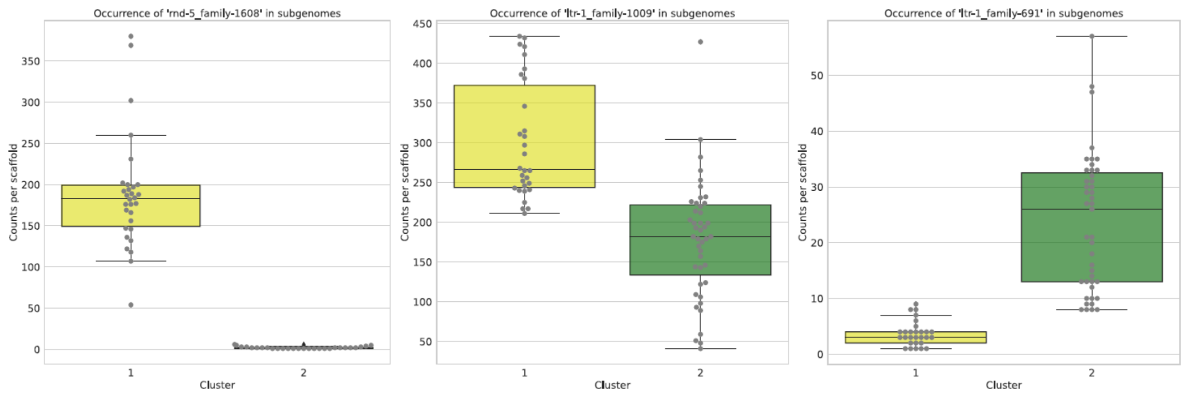


**Figure S10:** Repeat count of cluster specific and shared repeats between subgenomes. Occurrence of 3 distinct repeat families are shown as counts per scaffold (y-axis) that are divided over distinct clusters 1 and 2 (x-axis). Counts per scaffold are represented by grey dots. The repeat family identifier is indicated above each plot. The left panel shows the occurrence of an unclassified repeat family in cluster 1 specific scaffolds while absent in cluster 2 scaffolds. The right panel shows an unclassified cluster 2 specific repeat family. The unclassified repeat family in the middle graph is cluster unspecific.

**A**

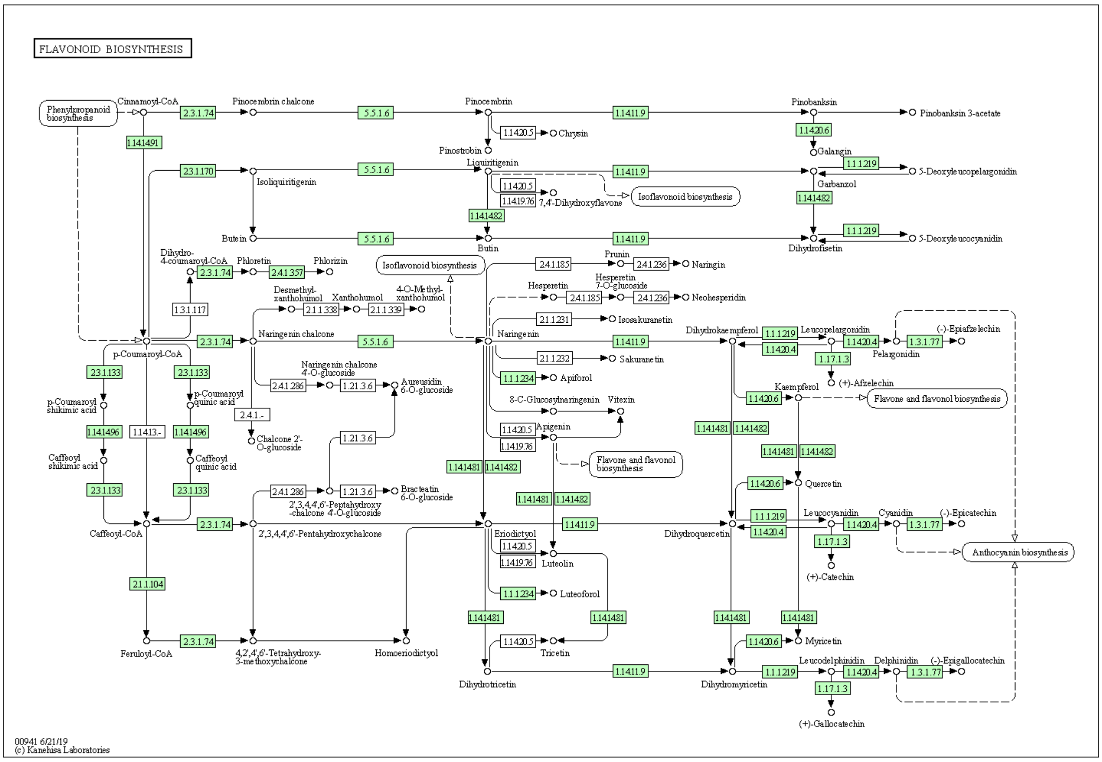


**B**

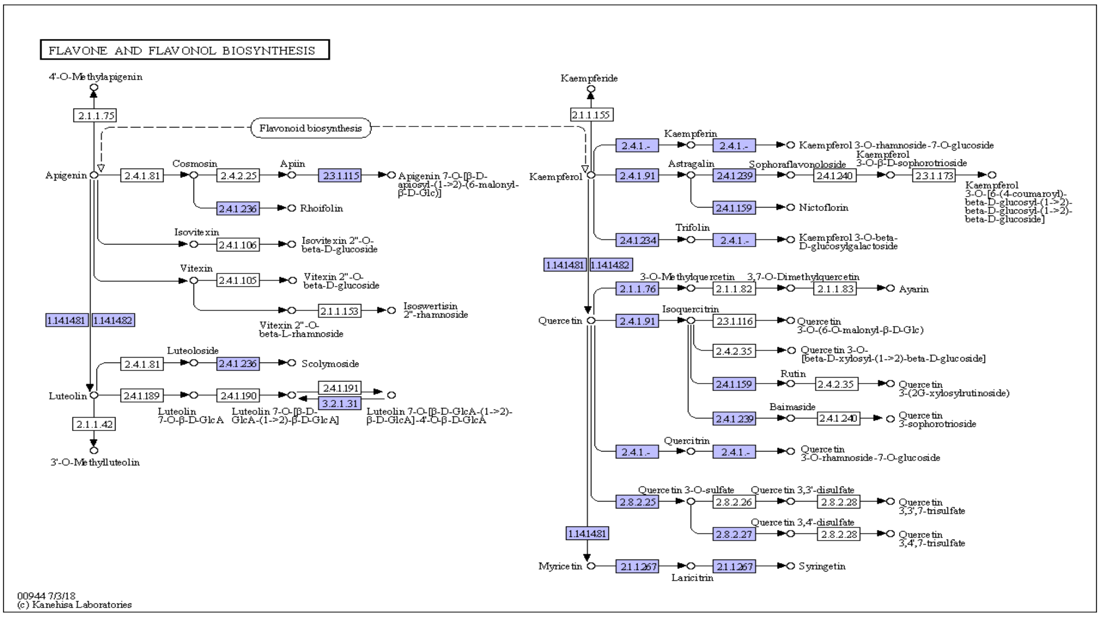


**Figure S11:** The flavonoid (A), and flavone and flavonol (B) KEGG bio-synthesis pathways in *Abelmoschus esculentus* (http://www.kegg.jp/kegg/kegg1.html). Putative okra enzyme coding genes for which a bi-directional best hit was found to enzymes pathways are shown with coloured EC identifiers.

### Supplementary tables

| Sample ID | Info | Rel. DNA amount | Flow Histogram |
|---|---|---|---|
| S-002 | Okra1 | | 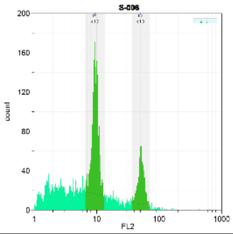 |
| S-003 | Agave | 15.90 | 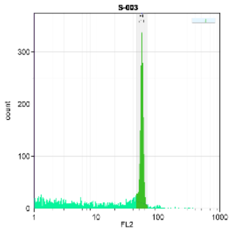 |
| S-004 | Okra1+Agave | 2.99 | 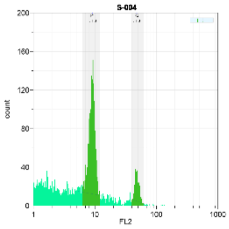 |
| S-006 | Okra2+Agave | 2.94 | 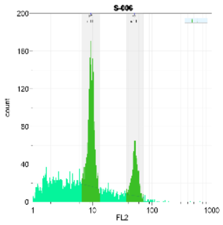 |
| S-007 | Okra3+Agave | 2.94 | 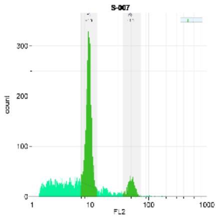 |
| S-008 | Okra4+Agave | 3.02 | 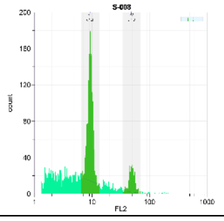 |
| S-009 | Okra5+Agave | 3.05 | 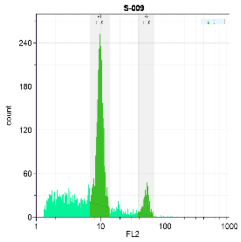 |
| **Average** | | 2.99 (±0.01) | |

**Table S1:** DNA amount of nuclei samples from okra root tip cells. DNA amount of okra replicate samples in picogram quantities was compared to a reference sample from *Agave Americana*. In the right column flow histograms of Okra samples and the Agave reference sample are shown. The count of observed nuclei in each histogram is depicted on the y-axis and is proportional to fluorescent intensity of each peak. The position of the peak along the x-axis is proportional to the relative DNA amount in each nuclei.

| Library type | Read length (bp) | raw yield (Gbp) | Read 1 >Q30 (bp) | Read 2 >Q30 (bp) |
|---|---|---|---|---|
| **10X PE** | 151 | 362 | 91.7 | 89.0 |
| **10X PE** | 151 | 318 | 92.9 | 90.4 |
| **10X PE** | 151 | 122 | 92.5 | 88.9 |
| **Library type** | **Av. polymerase read length (bp)** | **Subread length N50 (bp)** | **Longest subread** | |
| **Pacbio HiFi** | 411.39 | | 16,927 | |
| **Pacbio HiFi** | 516.25 | | 13,086 | |
| **Pacbio HiFi** | 472.40 | | 12,732 | |
| **Library type** | **Total molecules** | **Total length (bp)** | **Average length (kbp)** | **Molecule N50 (kbp)** |
| **BioNano unfiltered** | 77,407,575 | 4,887,375.86 | 63.138 | 90.175 |
| **BioNano filtered** | 5,500,210 | 1,184,887.6 | 215.426 | 206.88 |

**Table S2:** Genome sequencing and genome map data statistics. Linked read sequencing for three 10X Genomics libraries was performed using Illumina paired-end (PE) sequencing. Circular consensus sequencing (CCS) was performed for three Pacbio Hifi sequence libraries. Genome map data was produced for one BioNano DLE labelled library.

| Rank | No. hits | Hits | Family |
|---|---|---|---|
| **1** | 319 | Eukaryota *Gossypium hirsutum* | *Malvaceae* |
| **2** | 224 | Eukaryota *Theobroma cacao* | *Malvaceae* |
| **3** | 80 | Eukaryota *Gossypium raimondii* | *Malvaceae* |
| **4** | 72 | Eukaryota *Gossypium arboreum* | *Malvaceae* |
| **5** | 62 | Eukaryota *Durio zibethinus* | *Malvaceae* |
| **6** | 57 | No hits found | - |
| **7** | 29 | Eukaryota *Abelmoschus esculentus* | *Malvaceae* |
| **8** | 20 | Eukaryota *Hibuscus cannabinus* | *Malvaceae* |
| **9** | 14 | Eukaryota *Gossypioides kirkii* | *Malvaceae* |
| **10** | 11 | Eukaryota *Spondias tuberosa* | *Anacardiaceae* |

**Table S3:** BlastN screening statistics for Pacbio HiFi reads. Screening was performed against the NCBI nucleotide database, Readouts are indicated in counts of *Malvaceae* species specific and non-*Malvaceae* hits for a subset of 1000 HiFi reads.

| Rank | No. hits | Hits |
|---|---|---|
| **1** | 889 (88.9%) | No hits |
| **2** | 64 (6.4%) | *Malvaceae* mitochondria |
| **3** | 47 (4.7%) | Non-*Malvaceae* mitochondria |
| **Total** | 111 (11.1%) | mitochondria |
| **Rank** | **No. hits** | **Hits** |
| **1** | 919 (91.9%) | No hits |
| **2** | 44 (4.4%) | Non-*Malvaceae* chloroplast |
| **3** | 37 (3.7%) | *Malvaceae* chloroplast |
| **Total** | 81 (8.1%) | chloroplast |

**Table S4:** Pacbio sequence library contamination statistics for 1000 HiFi. Organelle content was determined using a BlastN screening against mitochondrial and chloroplast databases.

| Assembly statistic | Primary ctgs | Alternative ctgs | Hybrid scfds |
|---|---|---|---|
| **Ctgs/scfds** | 1417 | 526 | 124 |
| **Total length** | 1,223.6 Mb | 17.2 Mb | 1,194.5 Mb |
| **Median length** | 32.4 kb | 24.5 kb | 16.320 Mb |
| **Max length** | 25.1 Gb | 652 kb | 29.444 Mb |
| **Min length** | 13.3 kb | 10.3 kb | 125 kb |
| **N50 length** | 10.6 Gb | 30.5 kb | 18.929 Mb |
| **N50 index** | 43 | 126 | 27 |
| **N95 length** | 483 kb | 15.5 kb | 7.206 Mb |
| **N95 index** | 168 | 465 | 64 |
| **GC content** | 34.36% | 47.6% | 33.76% |

**Table S5:** NGS assembly and hybrid scaffolding statistics. Sequences were assembled using the Hifiasm assembler and scaffold with Bionano Genomics genome maps.

| Genome map statistic | Count |
|---|---|
| **Genome map count** | 216 |
| **Label density (/100kb)** | 15.94 |
| **Total genome map length (Mbp)** | 1,248.8 |
| **Genome map N50 (Mbp)** | 12.976 |
| **Total molecules aligned to genome maps** | 3,146,963 |
| **Fraction of molecules aligned** | 0.572 |
| **Effective coverage** | 374.509 |
| **Average confidence** | 21.8 |

**Table S6:** *De novo* genome map assembly statistics. Assembled molecules were mapped back to genome maps to estimate the effective coverage and average confidence the *de novo* assembly.

| **BUSCO class** | | **BUSCO distribution and topology** | | | | | | |
|---|---|---|---|---|---|---|---|---|
| **Copy nr** | **Count** | **1 ctg** | **Config.** | **2 ctgs** | **Configuration** | **3 ctgs** | **Configuration** | **4 ctgs** |
| **Single** | 284 | 284 | 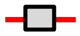 | 0 | | 0 | | 0 |
| **Duplicated** | 1150 | 1 | 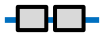 | 1149 | 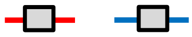 | 0 | | 0 |
| **Triplicated** | 843 | 0 | | 6 | 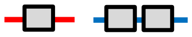 | 837 | 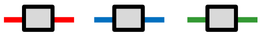 | 0 |
| **Quadruplicated** | 7 | 0 | | 1 | 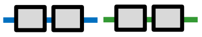 | 6 | 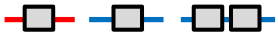 | 0 |
| **Quintuplicated** | 3 | 0 | | 0 | | 3 | 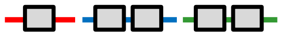 | 0 |
| **Sextuplicated** | 0 | 0 | | 0 | | 0 | | 0 |
| **Septuplicated** | 1 | 0 | | 0 | | 1 | 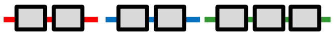 | 0 |
| **Total** | 2288 | 285 | | 1156 | | 847 | | 0 |

**Table S7:** BUSCO distribution and topology. BUSCO genes are classified according to their copy number in the genome. Distribution counts for ortholog gene copies have been indicated according to their position either on one, two or three contigs. Configuration of gene copies is depicted by a horizontal line representing a contig and superimposed small grey coloured boxes representing a gene copy.

| **Species** | **n/x** | **ploidy** | **identified** | **complete** | **duplicate** | **fragmented** | **missing** |
|---|---|---|---|---|---|---|---|
| A. thaliana | 2n=2x=10 | diploid | 98% | 98% | 17% | 0.5% | 2% |
| B. campestris | 2n=2x=18 | diploid | 99.4% | 99.1% | 14.7% | 0.3% | 0.6% |
| B. napus | 2n=4x=38 | allotetraploid | 97% | 97% | 90% | 1% | 3% |
| T. occidentale | 2n=2x=16 | diploid | 94% | 84% | 10% | 3% | 3% |
| T. pallescens | 2n=2x=16 | diploid | 94% | 83% | 11% | 2% | 4% |
| T. repens | 2n=4x=32 | allotetraploid | 94% | 24% | 57% | 3% | 6% |
| O. latifolia | 2n=2x=22 | diploid | 94.5% | 89% | 35% | 4.8% | 5.5% |
| B. amplexicaulis | 2n=6x=72 | hexaploid | 93.3% | 85% | 59% | 4.8% | 6.7% |
| S. lycopersicum | 2n=2x=24 | diploid | 96.8% | 96.4% | 1.0% | 0.3% | 2.1% |
| S. pennellii | 2n=2x=24 | diploid | 96.6% | 96.1% | 1.5% | 0.5% | 2.0% |
| S. lycopersicoides | 2n=2x=24 | diploid | 87.7% | 87.2% | 10.4% | 0,5% | 2.0% |

| | | | | | | | |
|---|---|---|---|---|---|---|---|
| S. chacoense M6 | 2n=2x=24 | diploid | 97% | 96% | 4.3% | 1% | 3.0% |
| S. tuberosum RH | 2n=2x=24 | diploid | 98.6% | 97% | 74.1% | 1.6% | 1.4% |
| S. tuberosum | 2n=4x=48 | autotetraploid | 94.2% | 85.7% | 9.5% | 8.5% | 5.7% |

| Tissue | Library type | Polymerase reads | | | Subreads | | Insert | |
|---|---|---|---|---|---|---|---|---|
| | | Size (Gbp) | Count | Mean length (bp) | Mean length (bp) | N50 (bp) | Mean length (bp) | N50 (bp) |
| Leaf | IsoSeq | 287.27 | 3,328,197 | 86,312 | 2,011 | 2,137 | 5,276 | 11,650 |
| Flower | IsoSeq | 323.48 | 6,676,796 | 48,448 | 1,723 | 1,774 | 4,536 | 8,917 |
| Pod | IsoSeq | 354.72 | 7,008,176 | 50,614 | 1,698 | 1,757 | 4,437 | 8,011 |

**Table S8:** Transcriptome sequencing statistics. Pacbio IsoSeq libraries were constructed for 3 different tissues as indicated.

| Sample | Type | Size (Gbp) | Mapper | Variant Caller | Map rate | SNPs |
|---|---|---|---|---|---|---|
| Haploid okra | IsoSeq | 20 | Minimap2 | GATK | 99.81% | 1109 |
| Commercial | IsoSeq | 1.6 | Minimap2 | GATK | 95.14% | 8127 |

**Table S9:** Transcriptome mapping to the okra reference genome and SNP calls.

| Superscaffold | configuration | #units | Start position | End position |
|---|---|---|---|---|
| 100090 | 18S-5.8S-28S | 24 | 13 | 289182 |
| 100111 | 18S-5.8S-28S | 54 | 8117 | 578985 |
| 100264 | 18S-5.8S-28S | 26 | 1 | 315695 |
| 67 | 18S-5.8S-28S | 32 | 2927 | 361934 |
| 74 | 18S-5.8S-28S | 5 | 20588136 | 20647257 |
| 8 | 18S-5.8S-28S | 8 | 1887219 | 2602099 |
| 28 | 18S-5.8S-28S | 11 | 8169258 | 8465035 |
| 34 | 5S | 8674 | 12382260 | 14843474 |

| | | | | |
|---|---|---|---|---|
| 34 | 5S | 71 | 14843474 | 18230209 |
| 44 | 5S | 16 | 8621481 | 8625812 |
| 62 | 5S | 852 | 890588 | 1182219 |

**Table S10:** Ribosomal gene clusters in the okra genome. Ribosomal gene clusters are characterized by unit configuration and number of tandemly arranged unit copies per superscaffold. Total lengths of clustered units that can be derived from the start and end position.
